# Environment-driven trends in fish larval abundance predict fishery recruitment in two temperate reef congeners: Mechanisms and implications for fishery recovery under a changing ocean

**DOI:** 10.1101/2023.10.11.561723

**Authors:** Erica T. Jarvis Mason, William Watson, Eric J. Ward, Andrew R. Thompson, Brice X. Semmens

**Affiliations:** Scripps Institution of Oceanography, University of California San Diego, California, 92037, USA; Southwest Fisheries Science Center, NOAA Fisheries, California, 92037, USA; Northwest Fisheries Science Center, NOAA Fisheries, California, 98112, USA

**Keywords:** Ecosystem-based management, data limited, species distribution model, fishery recovery, recruitment processes, fishery cohort strength, environmental indicators

## Abstract

Environmental and biological processes acting on fish larvae can drive fishery cohort strength, but predictive ability oftentimes falls short, and larval abundance is generally considered more useful as a proxy for spawning biomass. Under a changing ocean, studies that relate environmental covariates, larval abundance, and fishery recruitment are worthy of continued research, especially in data-limited contexts. We focus on a popular, recreational-only, multispecies saltwater bass fishery (genus *Paralabrax)* whose population status and recovery potential are uncertain. We used 54 years of ichthyoplankton data (1963-2016) and a species distribution model to 1) deconstruct species-specific standardized indices of larval abundance, 2) test these indices as indicators of adult stock status or predictors of future fishery recruitment, and 3) evaluate spatiotemporal trends in their population dynamics relative to environmental variables. Contrary to expectation, species-specific larval abundance predicted future catch, with recent elevated larval abundance suggesting imminent fishery recovery. Additionally, we identified strong relationships with environmental variables, thereby providing additional tools for predicting fishery recruitment and anticipating population change. Our findings paint a path forward for improving estimates of current and future fishery status under changing natural and anthropogenic influences and the incorporation of ecosystem considerations into fishery management.

## Introduction

An ecosystem approach to fishery management (Dolan et al. 2016) includes consideration of the many biological and environmental factors that have historically, and are currently, influencing fished populations. This, of course, also requires having information and resources to adequately evaluate the status of fish populations. Unfortunately, many small-scale fisheries lack such a robust approach, partly because data and resources are lacking. Given that the economic and ecological impacts of some marine recreational fisheries have rivaled those of commercial fisheries (Cooke & Cowx 2004, Lewin et al. 2019), and that overexploitation dampens the ecosystem resilience of fish populations (Perry et al. 2010, Ziegler et al. 2023), there is a clear need for research that leads to more robust population assessments for recreational-only and small-scale commercial fisheries.

One of the most popular and economically important recreational-only marine fisheries in California, USA, and perhaps even the world, is the southern California saltwater bass *Paralabrax* spp. fishery, which consists of Barred Sand Bass *P. nebulifer* (BSB), Kelp Bass *P. clathratus* (KB), and Spotted Sand Bass *P. maculatofasciatus* (SSB). This multi-species fishery has persisted since the early 20^th^ century, but catches have remained at historic lows since 2013 and are thought to primarily reflect a depressed population of BSB (a species targeted during spawning aggregations; Jarvis et al. 2010, 2014a, Erisman et al. 2011). Dramatic catch declines for BSB and KB began in 2005, following a period of increasing fishing mortality and population recruitment failure (Jarvis et al. 2014a). In contrast, SSB is almost entirely catch-and-release, limited to bays and estuaries, and presumed to have a healthier population. In 2013, regulations for the saltwater basses became more restrictive (Cal. Code Regs. Tit. 14, § 28.30; Jarvis et al. 2014a), but the fishery has not recovered, casting further uncertainty into the status and recovery potential of the BSB and KB populations. Adding to that uncertainty is the unknown impact that a warming ocean will have on these populations. Thus, research on the population dynamics of the saltwater basses may help with anticipating change and aid in the rebuilding and future management.

Differences in the geographic ranges, habitats, and reproductive strategies of the three saltwater basses suggest different sensitivities to fishing pressure and environmental conditions, and thus, differences in their resilience to climate change impacts. KB have occurred as far north as the cool temperate coast of the state of Washington, USA, while BSB and SSB have a maximum northern range extent that is approximately 1,000 km to the south, off the central California coast (Love and Passerelli 2020). The southernmost occurrence for KB is southern Baja California, Mexico, and for BSB and SSB, it is equatorward in the tropical waters of Acapulco, Mexico; SSB also have populations in the Gulf of California (Heemstra 1995). As a group, the saltwater basses have responded to decadal shifts in oceanographic conditions throughout the history of the fishery, being relatively more abundant during warmer ocean temperature regimes (Moser et al. 2001b, Hsieh et al. 2005, Jarvis et al. 2014a) and at one time, more common north of southern California (Hubbs 1948). Therefore, it is possible that northern latitudinal shifts in saltwater bass larvae abundances occur naturally, coinciding with decadal-scale oceanographic cycles and climate-driven increases in sea surface temperature (SST, Auth et al. 2018). Yet, at finer temporal scales, the saltwater basses may show differential population responses associated with seasonal or interannual oceanographic variability, i.e., upwelling, El Niño. Thus, understanding the relative role environmental variability has had on saltwater bass populations with respect to their vulnerability to harvest, i.e., catch-and-release versus aggregation-based, should help elucidate which species may be resilient to climate change impacts.

Southern California is uniquely data-rich in terms of long-term coastal oceanography data and thus represents an ideal testbed for the development of ecosystem-assessment methods. The California Cooperative Oceanic Fisheries Investigations (CalCOFI) is the world’s largest and longest-standing fisheries oceanography survey (Gallo et al. 2019) and since 1951 has conducted quarterly surveys to collect biological samples (e.g., fish larvae abundance, zooplankton biomass) and hydrographic data at discrete sampling locations throughout the Southern California Bight (SCB). Due to its long temporal coverage and high spatial resolution, the survey has enabled researchers to detect natural and anthropogenic influences on larval fish populations in southern California (Hsieh et al. 2009, Asch 2015, Thompson et al. 2017, 2022) and to assist resource managers in fishery assessments (Moser et al. 2001b, McClatchie 2014, Gallo et al. 2019). Combined with other oceanographic and fishery data streams, CalCOFI provides the unique opportunity to assess historical and recent spatiotemporal trends in saltwater bass populations. Indeed, interest in parsing the CalCOFI *Paralabrax* spp. larval time series to species funded research in the 1970s and 1980s to develop morphological and genetic tools for doing so (Butler et al. 1982, Graves et al. 1990), but successful species discrimination across larval stages was only recently made possible with the development of a robust, species-specific taxonomic key (Jarvis Mason et al. 2022).

Larval abundance has long been considered a proxy for spawning stock biomass (Hilborn & Walters 1992, Cowan & Shaw 2002). Indeed, CalCOFI fish larvae have been incorporated in west coast fishery stock assessments or used for monitoring status and trends (see Gallo et al. 2019 for examples). On the one hand, if we assume bass larval abundance is a function of the biomass of reproductive females in the population, then fishery managers are provided with an easy, reliable means of tracking the status and trends of adult stocks. On the other hand, larval abundance can be used to predict fishery recruitment in some fish (Murphy et al. 2018, Tripp et al. 2023); that is, since larval fish dynamics are so highly influenced by environmental variability, the processes affecting larval growth and mortality are thought to drive future fishery year class strength (Hjort 1914, Lasker 1984, 1987, Cushing 1990, Houde 2001). However, due to continued abiotic and biotic pressures on subsequent life stages, the accuracy of recruitment prediction from larval indices is generally considered to be better for species with lower ages of recruitment to fisheries (e.g., the juvenile or subadult stage, Bradford 1992, Cowan & Shaw 2002). Jarvis et al. (2014a) reported strong relationships between combined *Paralabrax* spp. larval abundance and species-specific fishery recruitment strength between 1997 and 2012, but it is unknown whether this relationship translates using species level larval abundance over a longer period.

Here, we unlock the CalCOFI *Paralabrax* spp. larval archive for the first time. In doing so, we intend to,

I. generate indices of southern California bass larvae abundance to improve species-specific estimates of population status and trends,
II. test our indices as indicators of adult stock status or predictors of future fishery recruitment, and
III. evaluate spatiotemporal trends in bass larval abundance relative to environmental variables and climate forcing (e.g., latitudinal shifts in abundance).

In the face of a changing ocean, the ability to identify potential environmental indicators of population status or future fishery recruitment, i.e., potential management action triggers, will be ever more important for guiding sustainable fishery management, especially in data-limited contexts, and may be a key determinate in the fate of this historical fishery.

## Methods

### Survey and study region

We focused on *Paralabrax* spp. larvae collected in CalCOFI oceanographic cruises conducted along the continental shelf off southern California, USA and Baja California, Mexico during July cruises from 1963 to 2016 (Fig. 1a, Table 1) because: 1) July corresponds to peak spawning (Walker et al. 1987), 2) *Paralabrax* spp. larvae occur nearshore relative to most other economically important groundfish in the SCB (McGowen 1993, Moser et al. 2001a), and, 3) at the time of this study, the CalCOFI ichthyoplankton archive was being updated in reverse chronological order to current taxonomic standards (1963 and 2016 were the most historical and recent years available, respectively). Two changes to the CalCOFI zooplankton sampling occurred during this period, 1) in 1969, the depth from which nets were obliquely towed increased from 140 m to 210 m (or from within 5 m of the seafloor in shallower waters), and 2) in 1978, the sampling gear changed from ring nets to paired bongo nets (Thompson et al. 2017). Survey coverage has varied spatially and temporally, with more consistent temporal coverage in the SCB (Fig. 1a) and additional, shallower, coastal stations added in 2004 (Gallo et al. 2019).

**Figure 1.**
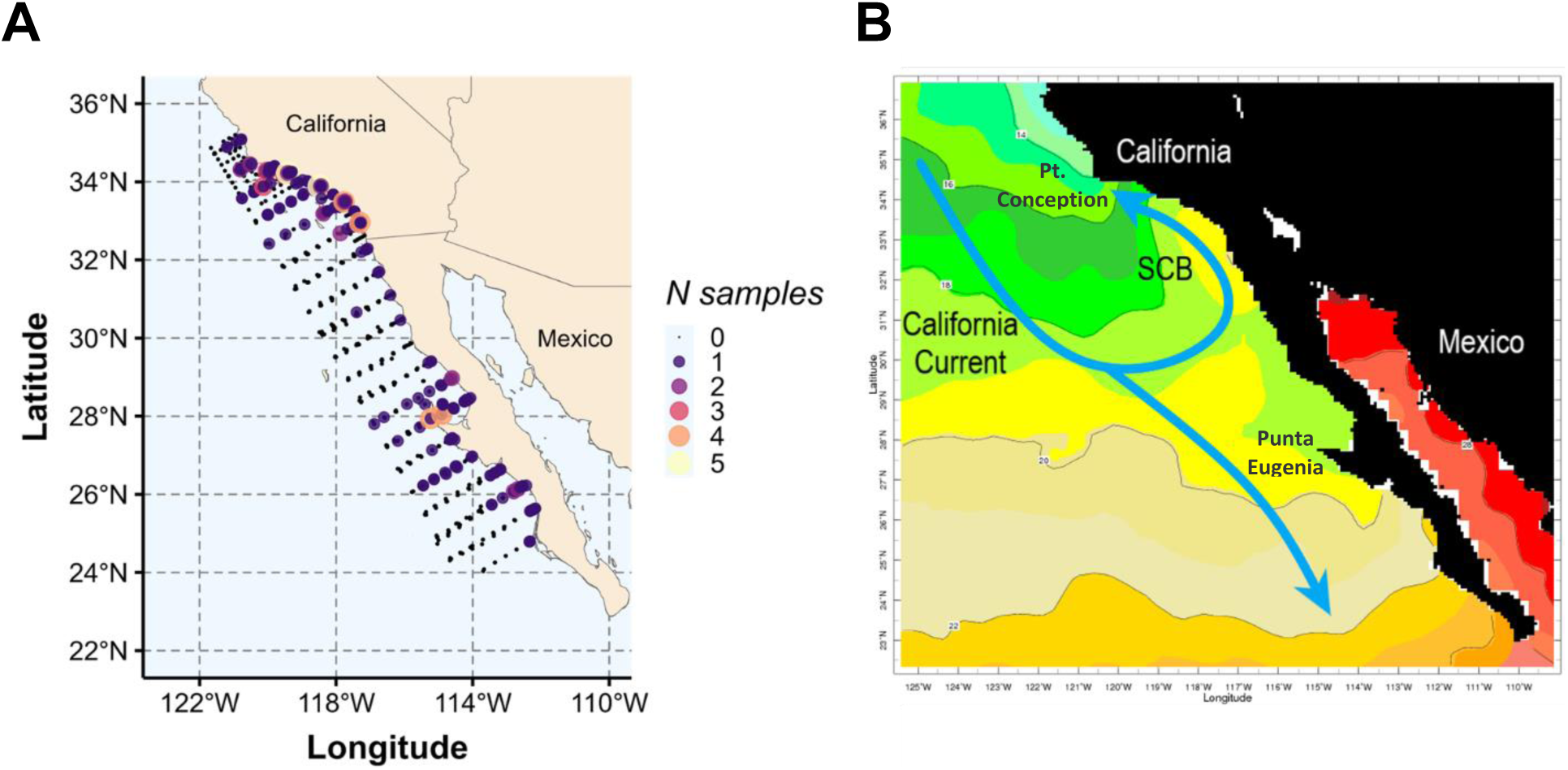
Maps of a) CalCOFI station locations surveyed along transect lines during July cruises from 1963 to 2016 and numbers of samples per station from which formalin preserved *Paralabrax* spp. larvae were identified to species, and b) representative sea surface temperature contours off southern California, USA and Baja California, Mexico in the month of July (NOAA NCDC OISST version2p1 AVHRR monthly sea surface temperature data). Blue arrows depict the directional flow of the California Current. CalCOFI = California Cooperative Oceanic Fisheries Investigations, SCB = Southern California Bight. Shaded temperature contours range from 14 °C in the north to 26 °C off the southern tip of Baja California and northward into the Gulf of California, Mexico.

**Table 1.**
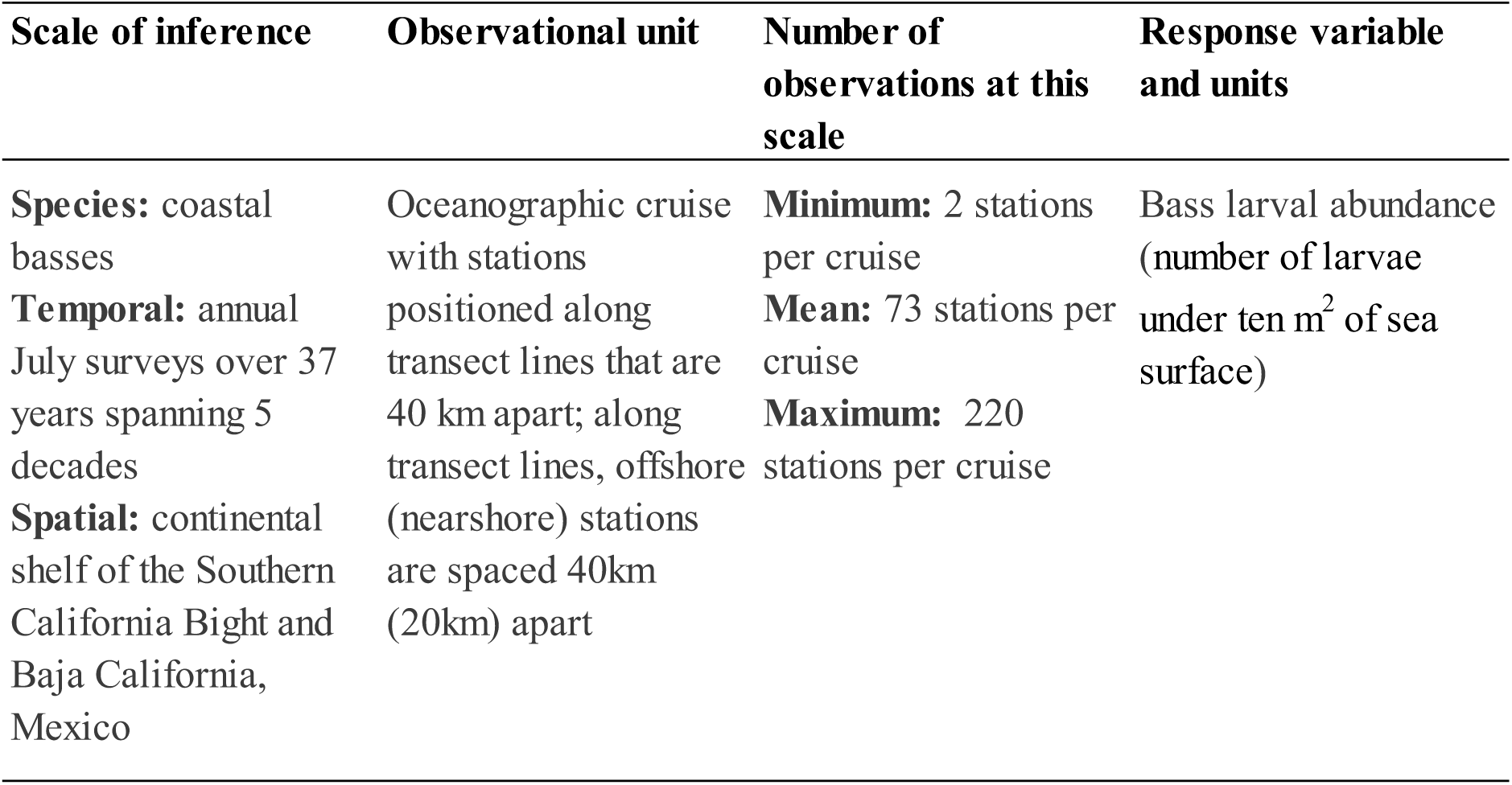
Temporal and spatial characteristics of California Cooperative Fisheries Investigations cruises selected for analyzing temporal and spatial trends in species-specific bass *Paralabrax* spp. larval abundance, 1963-2016.

| Scale of inference | Observational unit | Number of observations at this scale | Response variable and units |
| --- | --- | --- | --- |
| **Species:** coastal basses<br>**Temporal:** annual July surveys over 37 years spanning 5 decades<br>**Spatial:** continental shelf of the Southern California Bight and Baja California, Mexico | Oceanographic cruise with stations positioned along transect lines that are 40 km apart; along transect lines, offshore (nearshore) stations are spaced 40km (20km) apart | **Minimum:** 2 stations per cruise<br>**Mean:** 73 stations per cruise<br>**Maximum:** 220 stations per cruise | Bass larval abundance (number of larvae under ten m<sup>2</sup> of sea surface) |

The study region is part of the California Current Ecosystem, in which the offshore California Current (CC) brings cool, nutrient-rich waters equatorward from north of Pt. Conception, CA (34.2°N), before branching shoreward toward the U.S.A./Mexico border (32.2°N), and then poleward along the southern California coast, forming the counterclockwise southern California Eddy within the SCB in the summer months (Fig. 1b, Hickey 1993, McClatchie 2014). The southern branch of the CC continues to flow equatorward before veering westward offshore of Baja California Sur. Thus, the study region generally consists of cooler, temperate waters to the north and warmer, subtropical waters to the south (Fig. 1b, McClatchie 2014).

### Larval species identifications and data processing

Two taxonomists sorted and processed *Paralabrax* spp. larvae from archived CalCOFI cruise station vials. We assigned the larval stage of each larva according to notochord development and then assigned species using pigmentation patterns described in Jarvis Mason et al. (2022). We excluded yolk sac larvae as these could not be reliably identified, and we excluded *P. auroguttatus* (Goldspotted Sand Bass) because its population is rare north of Baja California Sur (Moser et al. 1996). We photographed all larvae and recorded the presence/absence of stage-specific morphological characters described in Jarvis Mason et al. (2022). For quality assurance, we selected a random, temporally stratified, subset of samples (10%) to be identified by both taxonomists. We reviewed and resolved all identifications with taxonomic disagreement. Four station vials contained hundreds of *Paralabrax* spp. larvae, and we subsampled at least 10% of the contents of these vials and then extrapolated the raw species counts based on the proportion of each species identified in the subsample, including any unidentified or yolk sac larvae. The two taxonomists regularly collaborated to resolve outstanding, difficult-to-identify larvae.

To characterize the overall sample of visually identified *Paralabrax* spp. larvae, we summarized the raw numbers of larvae by species, larval stage, and region (southern California, Baja California) and we plotted their geographic distributions. For subsequent analyses, we standardized species counts in each sample by the standard haul factor to obtain the number of larvae under ten m^2^ of sea surface (Thompson et al. 2017). We refer to this standardized raw count as larval abundance. There were 26 samples from 12 cruises in which larvae were so damaged as to not be identifiable to species or there was a discrepancy in the number of *Paralabrax* spp. larvae reported in the database versus the number observed in the sample vial (e.g., missing larvae). In this case, we first rechecked the original station vials and reference collection for the missing larvae, and if not found, then we estimated larval abundance (the standardized counts) for the sample based on the average proportion of each species standardized count at that station over the course of the entire study period.

### Standardized index of larval abundance

We used species distribution models (SDMs) to generate standardized indices of bass larval abundance in southern California. We did not include larval abundance data from Baja California because July CalCOFI did not survey the entire Baja California region after 1980. We used a geostatistical generalized mixed effects model framework that can incorporate spatial and spatiotemporal random fields (effects) to account for latent variables that may contribute to observations being correlated in space and time. The spatial random field captures latent spatial variation that is unchanging through time (e.g., distance to shore), while the spatiotemporal random field captures latent time varying spatial patterns (e.g., dynamic biological and oceanographic processes not captured by covariates in the model). This type of SDM can therefore account for unbalanced sampling effort and thus, is more precise than other SDM methods at estimating abundance (Thorson et al. 2015, Brodie et al. 2020). We fit SDMs with the R package `sdmTMB`, which estimates parameters of the Gaussian random fields using the Integrated Nested Laplace Approximation (INLA) and implements maximum marginal likelihood estimation with Template Model Builder (TMB, Anderson et al. 2022).

We modeled bass larval abundance as a function of variables that may affect “catchability” but not abundance (e.g., when and where you survey, gear type). For both species, we considered year, moon phase, day of year, hour, day/night by net type (to account for gear changes from ring to bongo nets, Thompson et al. 2017), and distance to the mainland coast. We did not consider the increase in tow depth because southern California *Paralabrax* spp. larvae are most abundant inshore of the 36 m depth contour (Lavenberg et al. 1986). We modeled bass larval abundance *Y* at location *s* and time *t* using a Tweedie observation error family (positive continuous density values that also contain zeros, Shono 2008) and a log link:

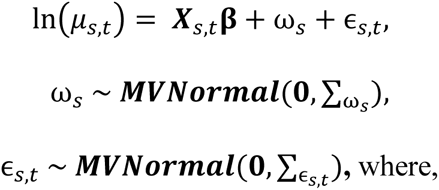

*μ_s,t_* represents mean abundance at location *s* and time *t*,

**X***_s,t_* is a vector of main effects, time varying effects, or spatially varying effects corresponding to location *s* and time *t*,

***β*** is a vector of estimated parameters, and

*ω_s_* and *ɛ_s,t_*, are the spatial and spatiotemporal random effects, respectively, assumed to be drawn from a multivariate normal distribution with covariance matrices ∑_ω,s_ and ∑_ɛs,t._ A user-specified triangulated mesh approximates the spatial component of the Gaussian random fields (Supplement S1, Anderson et al. 2022)

Given data gaps for some years, we could not model year as a fixed effect (factor). Instead, we modeled year effects and all other covariate effects with penalized smooth functions (penalized regression “P-splines”; Eilers and Marx 1996), similar to generalized additive models, GAMs, Wood 2017). We allowed the model to select the basis dimension, i.e., *k*, or wiggliness for each variable, except for the hour and continuous numeric moon phase covariates, in which we specified a cyclic smooth over 24-h and 4-w cycles, respectively. To account for potential day time net avoidance related to the gear change from ring to bongo nets, we specified day/night by net type as a factor smooth interaction where a separate smooth is created for each factor level. In cases of non-convergence, we tried simplifying variables to their linear form.

Using Restricted Maximum Likelihood (REML)-generated Akaike (AIC) values, we identified the best fit random effects model structure as an independent spatiotemporal random field across time steps (both species) and a spatial field estimated for BSB but not for KB. Using these species-specific random effects, we then compared model fit across different fixed effects (= main effects) with AIC (REML off). To allow for subsequent prediction of larval abundance in years without data, we specified missing years for model estimation of larval abundance. In the case of BSB, in which the inclusion of a smooth year effect was not supported, we incorporated year effects as a time-varying random walk (analogous to a dynamic linear model). We did not consider models that did not pass all model diagnostic checks (e.g., tests for convergence, positive definite Hessian matrix, large standard errors associated with model parameter estimates).

To generate an area-weighted standardized index of larval abundance, we used our model to predict abundance across a fine-scale grid covering the southern California CalCOFI survey region, with a grid cell resolution of 2 km x 2 km, focusing only on years when larvae were observed. This was intended to provide an index with less uncertainty across years. However, for relating larval abundance to catch data (see Relationship with fishery catch data below), in which missing years are not allowed, we generated the index of abundance based on the fine-scale prediction grid with missing years included, i.e., abundance was predicted for missing years.

### Spatiotemporal trends in larval abundance

To explore changes in larval abundance through time, we plotted spatiotemporal trends in larval abundance of both species using predicted annual abundances defined by a 2 km x 2 km resolution grid (exclusive of land and within the southern California CalCOFI survey area) and the parameter estimates generated from the index standardization model; missing years not included (Anderson et al. 2022). These trends included both fixed and random effects. We also separately plotted trends in the spatiotemporal random effects (*∊*_*s*,*t*_) to visually explore what patterns in the plots of fixed and random effects were being driven by latent spatiotemporal processes.

### Relationship with fishery catch data

For both species, we tested for a positive monotonic correlation between the index of larval abundance and spawning stock biomass, in which we used catch records from California Department of Fish and Wildlife (CDFW) Commercial Passenger Fishing Vessel (CPFV) logbooks (1975-2016) as a proxy for spawning stock biomass (Fig. S2, see Jarvis et al. 2014a for a description of catch data). The catch data represent the only index of saltwater bass abundance that matches the geographic extent of the CalCOFI larval abundance data. Typically, catch-per-unit-effort (CPUE) is considered an appropriate catch metric to use as an index of abundance because it accounts for changes in fishing effort over time. However, in this fishery, catch and CPUE directly correspond (Jarvis et al. 2014a) and species-specific CPUE at the trip level is not available prior to 1980. CPFV catch records reflect only fish kept (harvested catch) on commercial sportfishing vessels, the predominant fishing mode for BSB and KB (Jarvis et al. 2014a).

We considered only biologically plausible lags, i.e., catch cannot lead larvae, from zero to ten years and standardized all data to a mean of zero. We performed cross-correlation analysis using the R package `funtimes` (Lyubchich et al. 2023) in which a 95% confidence band provided guidance for interpreting correlations that may be influenced by the presence of autocorrelation in either or both datasets, i.e., coefficients falling within the band may be an artifact of non-stationarity.

We also explored the relationship between species-specific larval abundance estimates and estimates of *total* catch, i.e., estimates of harvest plus catch released alive or dead, by all fishing modes combined (e.g., CPFVs, private/rental boats, shore fishing). These data are available from the Marine Fisheries Statistical Survey (MRFSS, 1980-2003) and the California Recreational Fisheries Survey (CRFS, 2005-2016; PSMFC 2024). Estimates are derived from angler-intercept surveys of catch on a subset of fishing trips and extrapolated based on a phone survey of effort. MRFSS and CRFS are calculated using a different sampling design; however, since we were only concerned with monotonic relationships between larvae and catch, i.e., an increase in one variable predicts an increase (or decrease) in the other, we combined the time series and standardized values to a mean of zero (Fig. S2). Given that cross-correlation analysis does not allow for missing values, we imputed catch for four missing years in the total catch estimates for both species using the R package `mice` and the Predictive Mean Matching (PMM) method (van Buuren & Groothuis-Oudshoorn 2011).

We assumed a positive correlation between larvae and catch at a lag of 0 years to indicate a positive relationship between spawning stock biomass and larval abundance, where larval abundance is a function of the biomass of reproductive age females in the water, i.e., the higher the spawning stock biomass, the higher the larval abundance.

### Relationship with environmental variables

We also used the R package `sdmTMB` to model the influence of site-specific prey availability and temperature, as well as larger-scale oceanographic indices on bass larval abundance. For these models including environmental data, we specified a minimum distance cutoff of ten km in constructing the mesh.

Not all environmental variables of interest were available as far back as 1963, so we first modeled a smaller set of variables across the entire time series (1963 and 2016, Model 1) and then incorporated additional variables over the shorter period (1984 to 2016, Model 2). In addition, data available for modeling bass larval abundance was limited to those years for which a July CalCOFI cruise took place. For example, July CalCOFI cruises occurred triennially between the mid-1960s and mid-1990s, otherwise, July temperature and zooplankton data were available annually (Fig. 2). The larger scale environmental variables, North Pacific Gyre Oscillation (NPGO) index and Oceanic Niño Index (ONI), were also available annually, but the Biologically Effective Upwelled Transport Index (BEUTI), Coastal Upwelling Transport Index (CUTI), and isothermal layer depth (ILD) for southern California were only available from 1980 onward, while kelp canopy extent was available from 1984 onward (Fig. 2).

**Figure 2.**
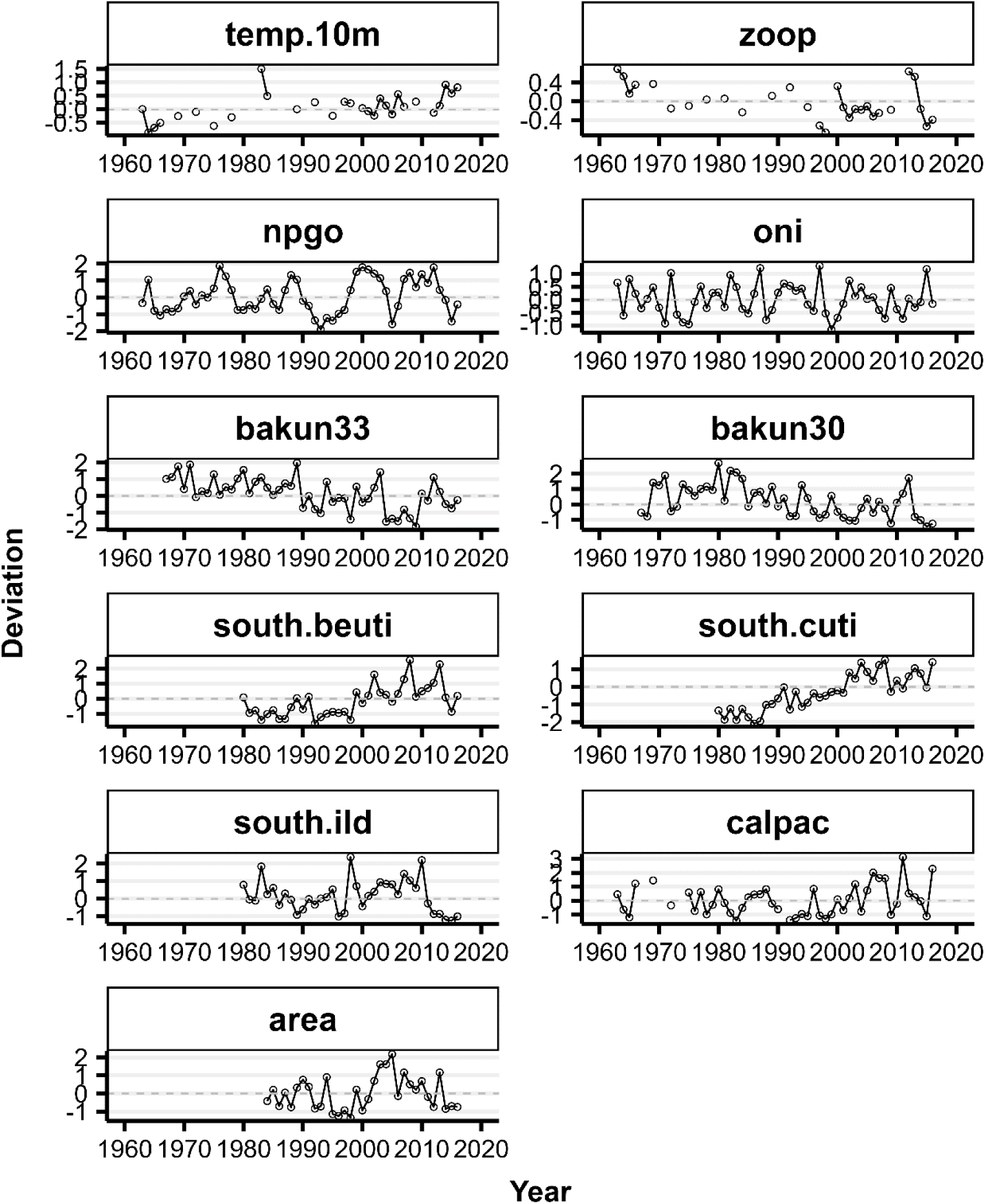
Temporal trends in regional and local environmental variables selected for modeling southern California, USA, *Paralabrax* spp. larval abundance. Temp.10m = CalCOFI ocean temperature averaged from the sea surface down to 10 m depth, zoop = CalCOFI zooplankton biomass, npgo = North Pacific Gyre Oscillation index for July, oni = Ocean Niño Index for June/July, bakun33 = Bakun upwelling index at 33°N, bakun30 = Bakun upwelling index at 30°N, south.beuti = Biologically Enhanced Upwelling Index, south.cuti = Coastal Upwelling Transport Index, south.ild = isothermal layer depth, calpac = spring biomass of the calanoid copepod, *Calanus pacificus*, area = areal canopy extent of Giant Kelp *Macrocystis pyrifera*. CalCOFI = California Cooperative Oceanic Fisheries Investigations.

For both species in Model 1, we included the station-specific CalCOFI temperature (°C, averaged across the upper ten meters) and square root transformed zooplankton biomass measured as zooplankton displacement volume (cm^3^/1000 m^3^ strained). We also included the NPGO index for the month of July and the ONI for June/July, and distance to the mainland coast (km, Fig. 2). The NPGO index is based on sea surface height variability throughout the north Pacific and tends to track regional nutrient fluctuations (or, mechanisms driving plankton ecosystem dynamics; Di Lorenzo et al. 2008). The ONI is based on surface temperature anomalies in the east-central equatorial Pacific and defines El Niño Southern Oscillation (ENSO) states (NOAA 2023).

For both species in Model 2, we included all variables from Model 1, and two additional variables: an upwelling index and the ILD in meters. We included upwelling because we wanted a more local measure of potential nutrient flux (versus the larger-scale NPGO), and we included ILD because spawning BSB orient to the thermocline during spawning season (McKinzie et al. 2014). In the study region, the ILD roughly corresponds to the mixed layer depth (MLD), which is the depth where temperature differs by 0.5 °C from the sea surface, i.e., we assume no salinity barrier layer between the MLD and ILD. We used a mean Bakun upwelling index (m^3^/sec) for June/July at latitude 33°N, where positive values indicate wind-driven offshore transport (Bakun 1973).

For both species, we considered another candidate Model 2 where we replaced the Bakun index with the BEUTI and CUTI specific to the southern California region, which provide a measure of nitrate flux through the mixed layer and coastal vertical transport, respectively (Jacox et al. 2018a, data from Hunsicker et al. 2022). Positive BEUTI values indicate nitrate flux into the mixed layer and positive CUTI values indicate coastal upwelling. For BSB, we additionally substituted upwelling in the SCB with upwelling off northern Baja California (Bakun index at latitude 30°N) because older recruits (∼ 2-y) showed a negative correlation with upwelling south of the SCB at a 2-y lag (Jarvis et al. 2004). For KB, we also considered a model that substituted upwelling in the SCB with kelp canopy extent because Giant Kelp *Macrocystis pyrifera* is important adult habitat (White & Caselle 2008). We obtained regional areal kelp canopy data by year from kelpwatch.org. Additionally, for both bass species, another candidate model considered (in place of zooplankton biomass) biomass of the calanoid copepod, *Calanus pacifus*, in spring CalCOFI cruises as a potential measure of larval pre-conditioning (Fennie et al. 2023, Swalethorp et al. 2023, Walsh 2023).

We used AIC to identify the most parsimonious among the Model 1 and Model 2 candidate models for each species. We ensured these species-specific candidate models were compared using the same random effects structure. We visually explored the conditional effects of important explanatory variables using the R package visreg (Breheny & Burchett 2019).

For all models, we standardized non-index variables to a mean of zero. We checked for multicollinearity among model covariates using the R packages corrplot (Wei & Simko 2021) and performance (Lüdecke et al. 2021). Like with other mixed models, our spatiotemporal model in sdmTMB did not permit inclusion of NA values, so prior to analysis we used the R package gstat (Pebesma 2004) to fill in missing within-cruise CalCOFI station data with the inverse distance-weighted interpolation method..

### Latitudinal shifts in larval abundance

To detect potential latitudinal shifts in bass larval abundance over time, we used larval abundance predictions from the standardized index of abundance models using only years with observed larvae and a species distribution function estimator within `sdmTMB` to calculate temporal trends in the center of gravity (COG) for both species (Thorson et al. 2016, Anderson et al. 2022). For larvae of both species, we plotted changes in the mean COG, grouped by decade and the mean July SST for each decade. We obtained July SSTs off Pt. Dume, CA, which is approximately centrally located along the southern California coast (Carter et al. 2022).

### Data and Code Availability

Data and code pertaining to the sdmTMB models and cross correlation analysis are available online in a GitHub repository: https://github.com/ETJarvisMason/xxx *(repository publicly available upon acceptance)*. We performed all analyses in R 4.3.2 (R-Core-Team 2023).

## Results

We examined a total of 1,267 “*Paralabrax”* spp. larvae sorted from original station vials (143 samples), of which 1,118 larvae could be positively identified; most (60.2%) were collected off Baja California, Mexico (Fig. S3). Larvae we observed in samples from southern California, USA, were 64.6% KB, 31.3% BSB, and 4.1% SSB. In samples taken off Baja California, numbers were highest for BSB (76.5%), followed by KB (16.5%) and SSB (7.0%). We encountered preflexion larvae more frequently across species and region; flexion and postflexion larvae were least common (Fig. S3). By station, KB were relatively more common offshore, while BSB and SSB were restricted to nearshore stations, with the highest numbers on average off Baja California Sur, south of Punta Eugenia (Fig. S4). SSB larvae in the CalCOFI time series were rare, particularly in southern California, indicating CalCOFI is not a representative survey for SSB larvae in this region (Fig. S3, S4).

There were four sub-sampled stations with high numbers of larvae that occurred off Baja California in the 1960s. Extrapolated species counts for these stations increased the overall raw larval count by an additional 1,716 larvae comprised of 93.0% BSB, 5.6% SSB, and 1.2% KB; we did not further extrapolate the counts by larval stage. The standardized raw counts of species-specific larval abundance in southern California indicated KB larvae encountered in CalCOFI bongo nets was generally higher year-to-year than that of BSB larvae (Fig. S5). Off Baja California, peaks in the standardized raw counts of bass taken in CalCOFI July samples occurred in the early 1960s and consisted primarily of BSB, while low counts from 1989 onward were representative of a single CalCOFI station off the very northern part of Baja California (Fig. S5), in which line 93.4 was the only Baja California CalCOFI line surveyed in those years. Cruises in 1981 and 1984 did not extend south beyond Punta Eugenia, Baja California (Fig. 1).

### Temporal trends in larval abundance

The most parsimonious index standardization models included BSB larval abundance as a time-varying function of distance to mainland coast (smoothed), incorporated all random effects, including an independent spatiotemporal field across years (“iid”); the resulting index for KB included larval abundance as a function of year (smoothed) with an independent spatiotemporal field across years and no spatial random effect (Table 2). Temporal trends in the standardized index of larval abundance for both species showed similar sporadic larval pulses in 1981, 1989, 2012, and 2014 (Fig. 3). Compared to BSB, KB larval abundance estimates were generally higher overall and between strong larval pulses. Larval abundance for both species was low between the late 1990s and early 2000s. On the natural log scale, we observed BSB larval abundance fluctuated more dramatically than KB larval abundance and declined steadily from the mid-1990s through the early 2000s before increasing again through the mid-2010s (Fig. 3a). In contrast, KB larval abundance fluctuated less than BSB larval abundance and showed a steady decline beginning in the early 1990s through the early 2000s before increasing again into the 2010s to levels similarly observed in the early 1980s (Fig. 3b).

**Figure 3.**
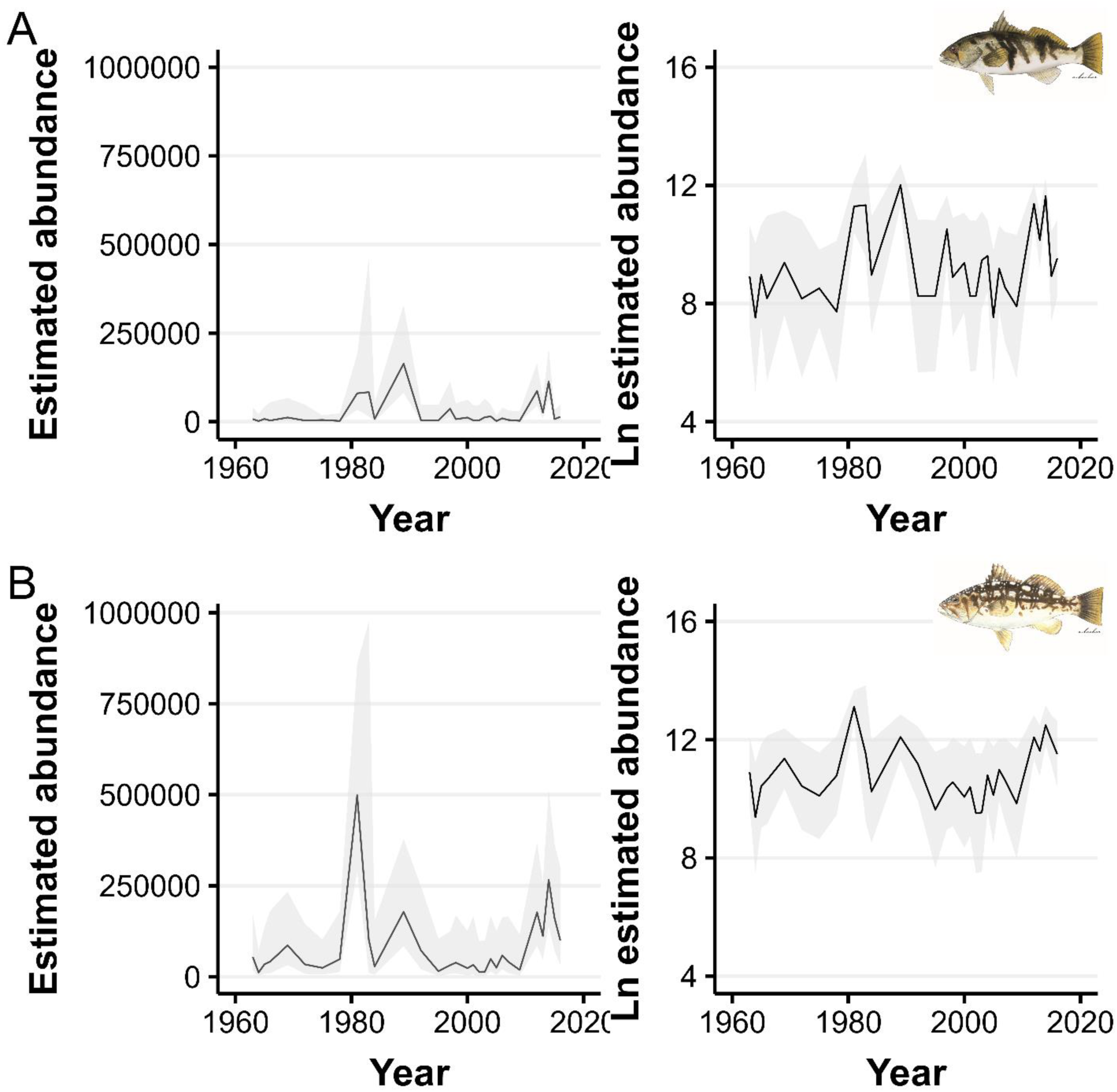
Standardized indices of larval abundance in normal space (left panels) and link space ln(abun), right panels) for a) Barred Sand Bass *Paralabrax nebulifer* and b) and Kelp Bass *P. clathratus* in southern California, USA, 1963-2016. Shaded ribbons denote 95% confidence intervals.

**Table 2.** Results comparison of candidate species distribution models* used to generate a standardized index of larval abundance for Barred Sand Bass *Paralabrax nebulifer* and Kelp Bass *P. clathratus* in southern California, USA, 1963-2016.

| Barred Sand Bass |  |  |  |  |  |
| --- | --- | --- | --- | --- | --- |
| Formula | Time-varying | Spatial | Spatio-temporal | REML AIC | AIC |
| ~ **0 + dist** | **y** | **on** | **iid** | **817** | **818** |
| ~ 0 + dist | y | off | iid | 828 | -- |
| ~ 0 + dist + day | y | on | iid | -- | 818 |
| ~ 0 + dist + s(moon, bs = "cc", k=4) | y | on | iid | -- | 820 |
| Kelp Bass |  |  |  |  |  |
| Formula | Time-varying | Spatial | Spatio-temporal | REML AIC | AIC |
| ~ **s(year)** | **n** | **off** | **iid** | **--** | **1484** |
| ~ s(year) + dist | n | off | iid | -- | 1527 |
| ~ s(year) + moon | n | off | iid | -- | 1486 |

\*Main effects are listed in the formula; random effects are spatial and/or spatiotemporal. Time-varying = main effects vary across years as a random walk, iid = independent and identically distributed across years, REML = restricted maximum likelihood, AIC = Akaike information criteria, dist = distance to mainland coast, day = day of year, moon = continuous numerical moon phase, bs = basis dimension, "cc" = cyclic.

### Spatiotemporal trends in larval abundance

Predicted bass larval abundance off southern California also showed geographic differences through time. BSB larval abundance was generally minimal throughout much of the SCB, with the highest densities occurring nearshore and primarily along the central and southern coast (Fig. 4). In contrast, KB larval abundance was relatively higher throughout the SCB, with hotspots occurring primarily at the northern Channel Islands (Fig. 5, see ‘2’ in Fig. S1 for location reference). KB larval abundance was particularly low throughout the SCB between 1995 and 2005. Spatiotemporal random effects plots for BSB and KB also showed different patterns through time (Fig. S6, S7), in which deviations from the fixed effects predictions varied depending on species, location, and year; KB tended to show more spatiotemporal structuring than BSB.

**Figure 4.**
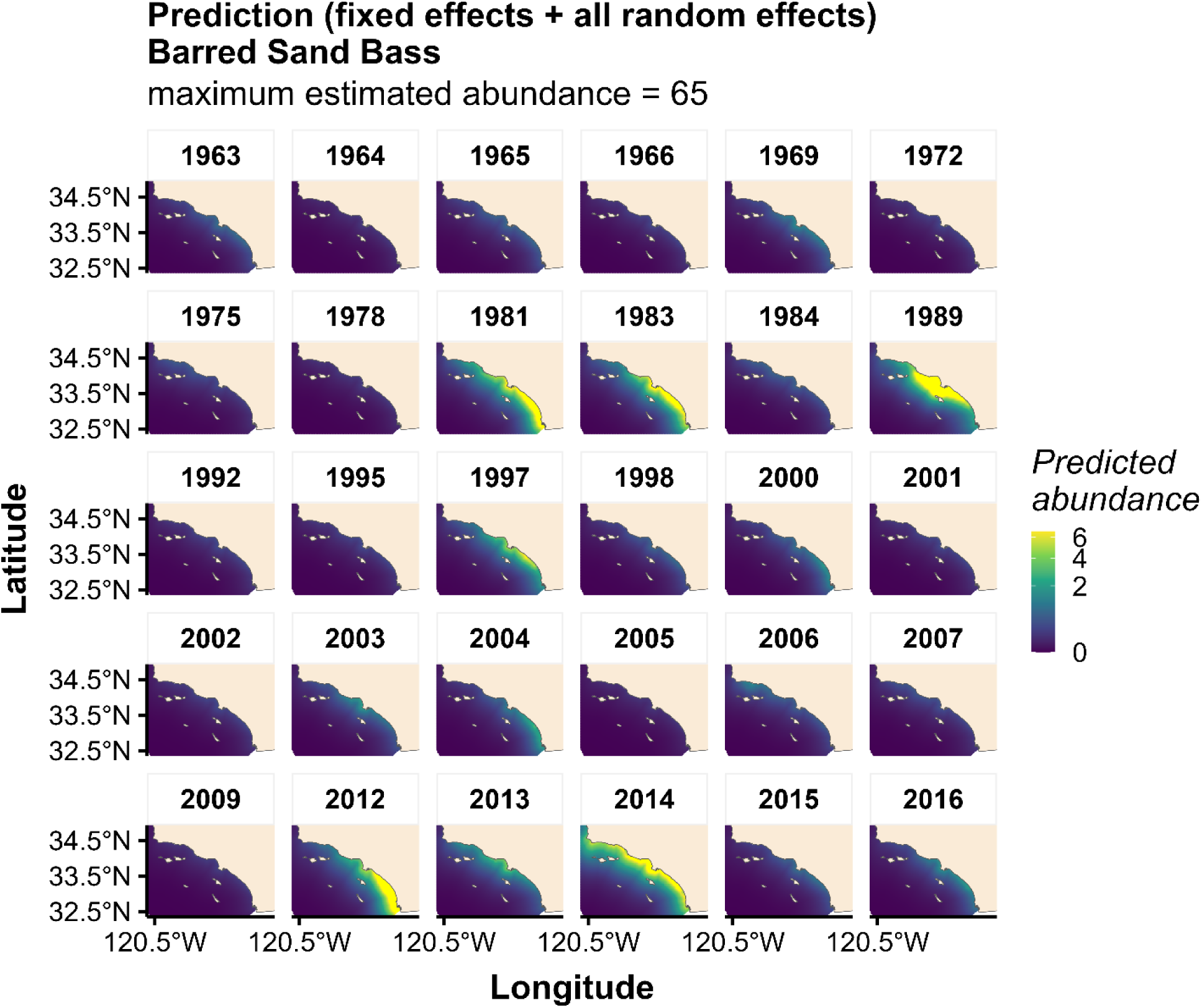
Predicted Barred Sand Bass *Paralabrax nebulifer* larval abundance in southern California, USA, by year, 1963-2016. Scale bar denotes abundance on the natural log scale. The legend color ramp ranging from light to dark denotes higher and lower abundance, respectively.

**Figure 5.**
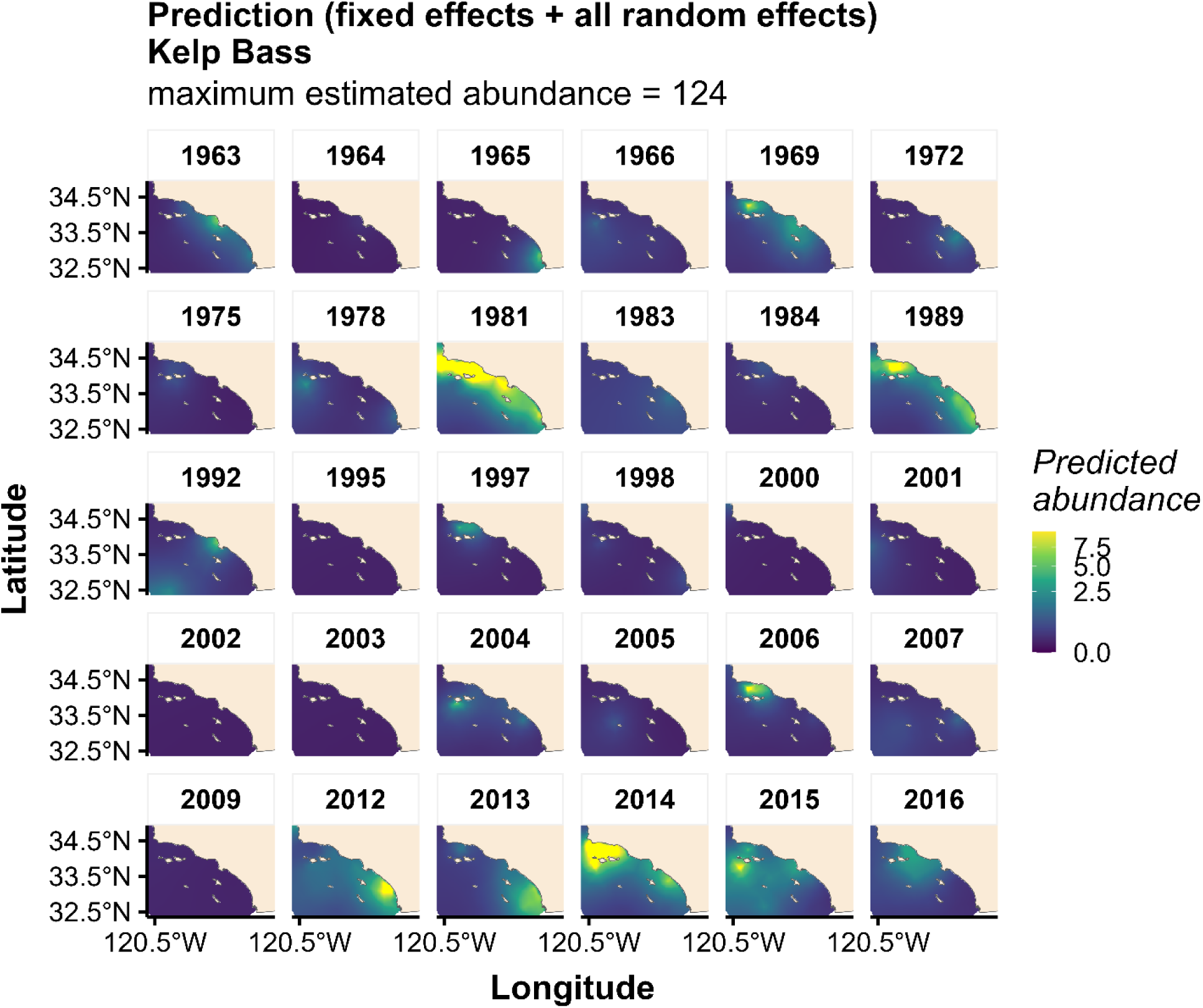
Predicted Kelp Bass *Paralabrax clathratus* larval abundance in southern California, USA, by year, 1963-2016. Scale bar denotes abundance on the natural log scale. The legend color ramp ranging from light to dark denotes higher and lower abundance, respectively.

### Relationship with fishery catch data

We found no relationship at a lag of zero between bass larval abundance estimates and fishery catch data (Fig. 6, see Fig. S8 for index estimated for all years). In contrast, for both species, we found marginal to strong lagged relationships between larval abundance and both sources of catch data. For example, the highest Spearman correlation coefficients between BSB larval abundance and total catch estimates occurred at 6-and 7-yr lags, corresponding to the age of most fishery recruits (Fig. 6a). We found a similar trend with the BSB harvest data, in which the highest correlations indicated larval abundance led catch by six to ten years. For KB, there were moderate to strong correlations between larval abundance and total catch estimates from 4-to 10-yr lags, corresponding to a higher catch-and-release rate across sizes for KB; KB larval abundance showed marginal correlations with harvest at 4-, to 6-yr lags and was strongly correlated with harvest at 7-to 10-yr lags (Fig. 6b).

**Figure 6.**
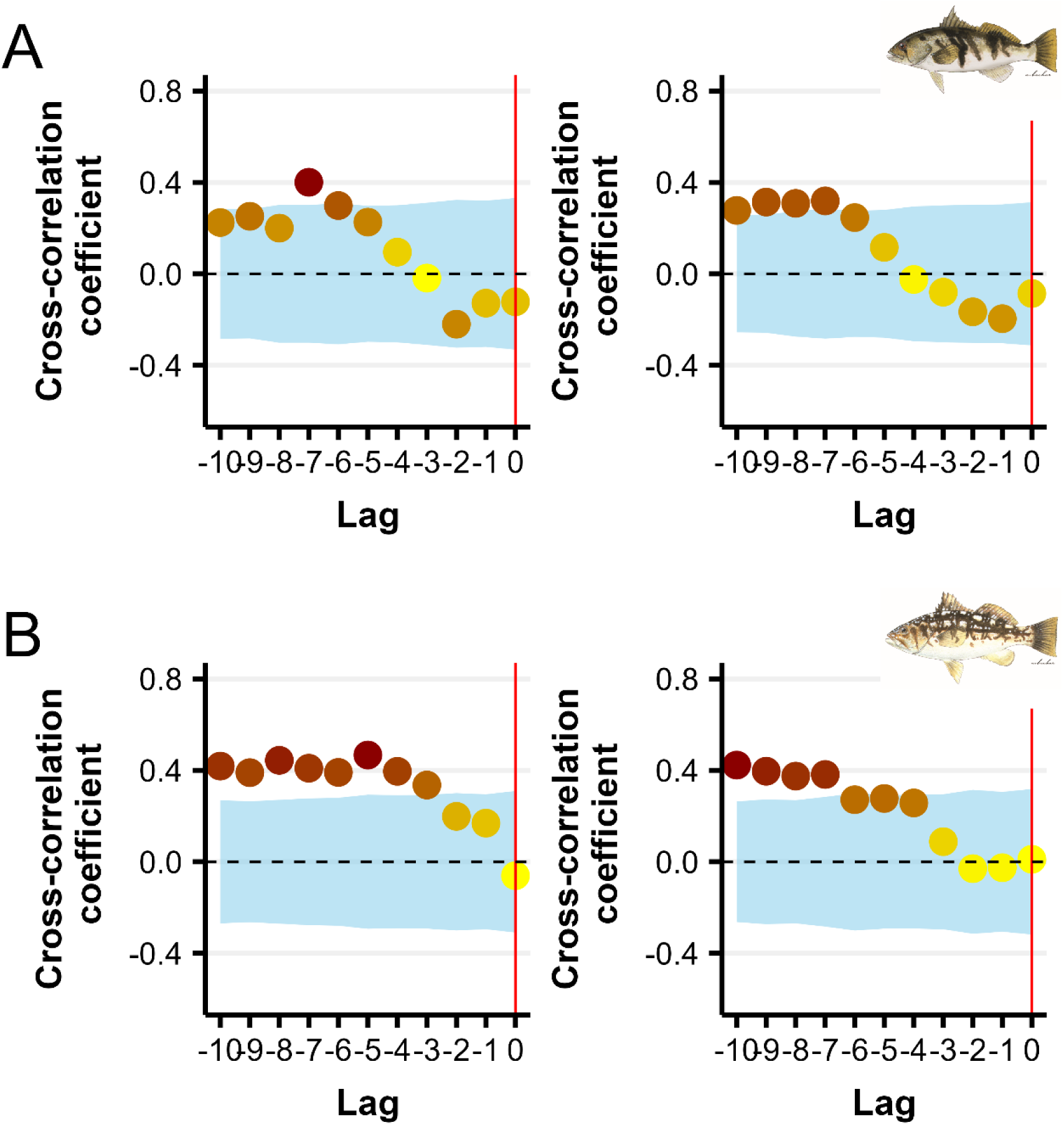
Cross-correlation Spearman coefficients between larval abundance and total catch estimates (left panels, includes estimates of fish kept and released across all fishing modes) and total numbers of reported harvest on CPFVs (right panels) across yearly lags for a) Barred Sand Bass *Paralabrax nebulifer* and b) and Kelp Bass *P. clathratus* in southern California, USA. CPFVs = Commercial Passenger Fishing Vessels. Negative lags represent larvae leading catch. Shaded ribbon denotes the 95% confidence band. Gradation in shading of points depicts the strength of the correlation, ranging from zero or weak (light) to marginal/moderate or strong (darker).

### Relationship with environmental variables

Among the 66 combinations of environmental variables, most correlations were weak (< 0.20, Fig. S9). Despite three relatively strong correlations; one between the BEUTI and the CUTI (0.71), one between the BEUTI and the NPGO (0.54) and one between the Bakun index 33°N and the NPGO (0.46), all model checks for multicollinearity produced “Low Correlation” results, with most covariate variance inflation factor values close to 1 and none greater than 4.

In southern California, over the entire study period, 1963 to 2016, zooplankton biomass and CalCOFI station-specific temperature (mean from surface to 10m) were positive predictors of bass larval abundance for both species (Fig. 7, S10, S11). There was also a strong negative relationship with distance to mainland coast (higher larval abundance nearshore; Fig. 7, S10, S11). The influence of temperature and distance to mainland coast was stronger for BSB than KB (Fig. 7).

**Figure 7.**
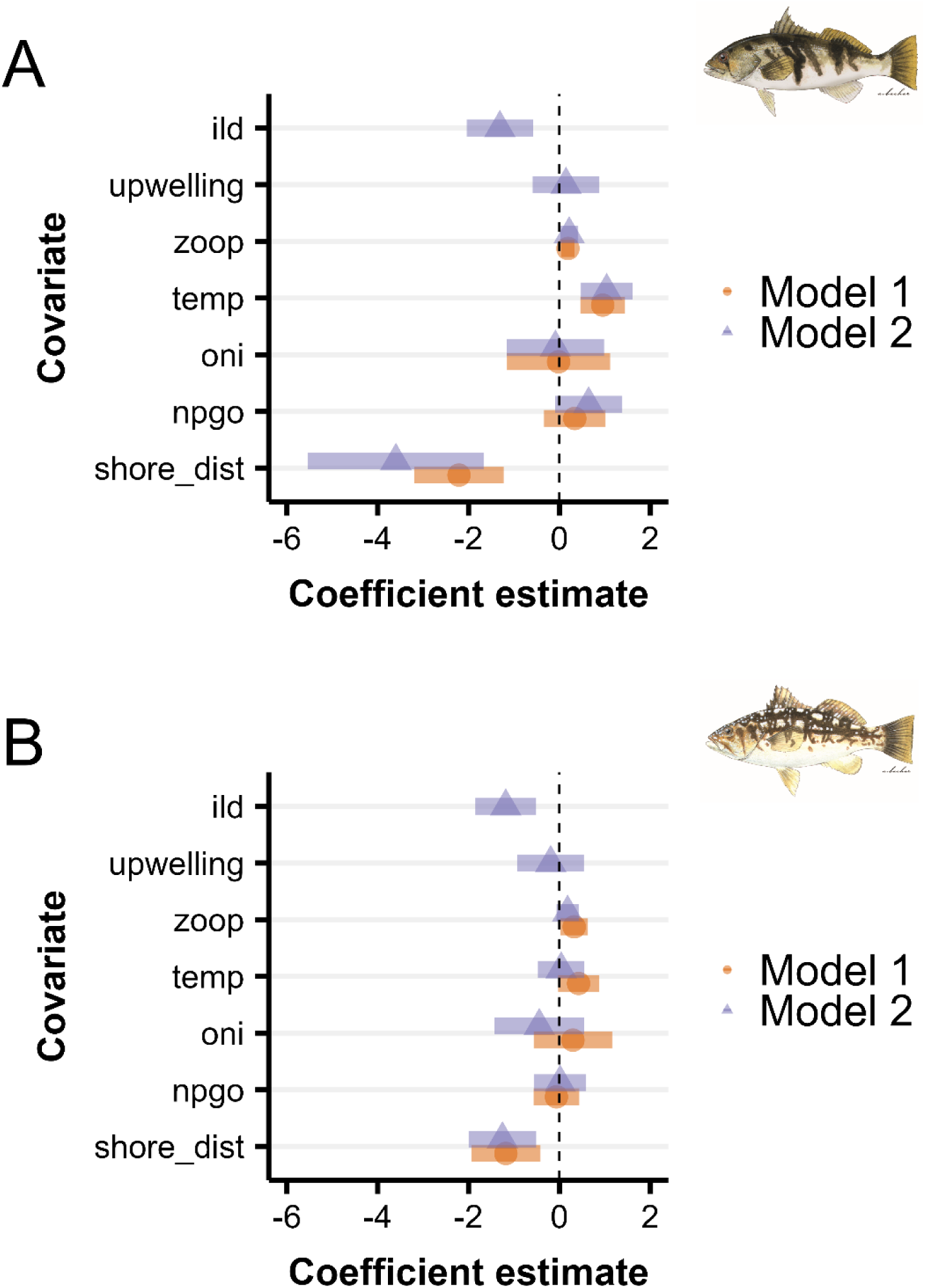
Model 1 coefficient estimates of the generalized linear mixed effects model depicting the relationships between bass larval abundance and environmental covariates from 1963 to 2016 (Model 1) and from 1984 to 2016 (Model 2) for a) Barred Sand Bass *Paralabrax nebulifer* and b) and Kelp Bass *P. clathratus* in southern California, USA. ild = isothermal layer depth, upwelling = Bakun upwelling index at 33°N, zoop = CalCOFI zooplankton biomass, temp = CalCOFI ocean temperature averaged over upper 10 m, oni = Ocean Niño Index for June/July, npgo = North Pacific Gyre Oscillation index for July, shore_dist = distance from survey station to mainland coast. Model coefficients greater (less) than one reflect positive (negative) relationships. Lines depict 95% confidence intervals. CalCOFI = California Cooperative Oceanic Fisheries Investigations.

Over the shorter study period, 1984 to 2016, representing Model 2, the ILD was strongly negatively related to bass larval abundance (high bass larval abundance with shallower ILD; Fig. 7, S12, S13). Of the different upwelling indices considered (Bakun, BEUTI, CUTI), models were essentially equally parsimonious, with only slightly lower AIC values for models with the Bakun index (Table 3); however, none of the upwelling indices showed a strong relationship to bass larval abundance. The positive relationship with zooplankton biomass and negative relationship with distance to mainland coast remained important for both species during the shorter period, while for BSB, temperature also remained important and the NPGO became more important (Fig. 7, S12). S13).

**Table 3.** Results comparison of candidate species distribution models* used to measure the effects of environmental influence on larval abundance for Barred Sand Bass *Paralabrax nebulifer* and Kelp Bass *P. clathratus* in southern California, USA during two time periods (Model 1, 1963-2016; Model 2, 1984-2016). Bold represents top model.

| Barred Sand Bass |  |  |  |  |  |
| --- | --- | --- | --- | --- | --- |
| Formula | Spatial | Spatio-temporal | Passed sanity check? | REML AIC | AIC |
| Model 1, 1963-2016 |  |  |  |  |  |
| ~ **dist_km + temp.10m + npgo + oni + zoop** | **off** | **iid** | **y** | **708** | **700** |
| ~ dist_km + temp.10m + npgo + oni + zoop | off | off | y | 766 | -- |
| ~ dist_km + temp.10m + npgo + oni + calpac | off | iid | y | -- | 749 |
| Model 2, 1984-2016 |  |  |  |  |  |
| ~ **dist_km + temp.10m + npgo + oni + zoop + bakun30 + south.ild** | **off** | **iid** | **y** | **--** | **576** |
| ~ dist_km + temp.10m + npgo + oni + zoop + bakun33 + south.ild | off | iid | y | -- | 578 |
| ~ dist_km + temp.10m + npgo + oni + zoop + south.beuti + south.cuti + south.ild | off | iid | y | -- | 578 |
| ~ dist_km + temp.10m + npgo + oni + calpac + bakun30 + south.ild | off | iid | y | -- | 619 |
| Kelp Bass |  |  |  |  |  |
| Formula | Spatial | Spatio-temporal | Passed sanity check? | REML AIC | AIC |
| Model 1, 1963-2016 |  |  |  |  |  |
| ~ **dist_km + temp.10m + npgo + oni + zoop** | **off** | **iid** | **y** | **--** | **1270** |
| ~ dist_km + temp.10m + npgo + calpac + oni | off | iid | y | -- | 1303 |
| Model 2, 1984-2016 |  |  |  |  |  |
| ~ **dist_km + temp.10m + npgo + oni + zoop + bakun33 + south.ild** | **off** | **iid** | **y** | **--** | **939** |
| ~ dist_km + temp.10m + npgo + oni + zoop + bakun30 + south.ild | off | iid | y | -- | 940 |
| ~ dist_km + temp.10m + npgo + oni + zoop + kelp + south.ild | off | iid | y | -- | 940 |
| ~ dist_km + temp.10m + npgo + oni + zoop + south.beuti + south.cuti + south.ild | off | iid | y | -- | 940 |
| ~ dist_km + temp.10m + npgo + oni + calpac + bakun33 + south.ild | off | iid | y | -- | 973 |

\* Main effects are listed in the formula; random effects are spatial and/or spatiotemporal. iid = independent and identically distributed across years, REML = restricted maximum likelihood, AIC = Akaike information criteria, dist\_km = distance to mainland coast, temp.10m = average CalCOFI sea surface temperature to 10 m depth, npgo = North Pacific Gyre Oscillation index for July, zoop = CalCOFI zooplankton biomass, oni = Oceanic Niño Index for June/July, calpac = CalCOFI spring biomass of the calanoid copepod, *Calanus pacificus*, bakun33 = Bakun upwelling index at 33° N, south.ild = isothermal layer depth, bakun30 = Bakun upwelling index at 30° N, south.beuti = Biologically Enhanced Upwelling Index, south.cuti = Coastal Upwelling Transport Index, kelp = areal canopy extent of Giant Kelp *Macrocystis pyrifera*, CalCOFI = California Cooperative Oceanic Fisheries Investigations.

### Latitudinal shifts in bass larval abundance

Bass larval abundance center of gravity within the SCB remained remarkably stable between the 1960s and 2010s (Fig. 8). The larval BSB center of gravity remained centrally located along the mainland coast, fluctuating off Santa Monica Bay to the north and Huntington Beach to the south (Fig. 8a). In contrast, the larval KB center of gravity was shifted offshore between the northern and southern Channel Islands (Fig. 8b). We observed no consistent northward or southward directional movement in the center of gravity for either bass species either by decade or mean decadal July sea surface temperature.

**Figure 8.**
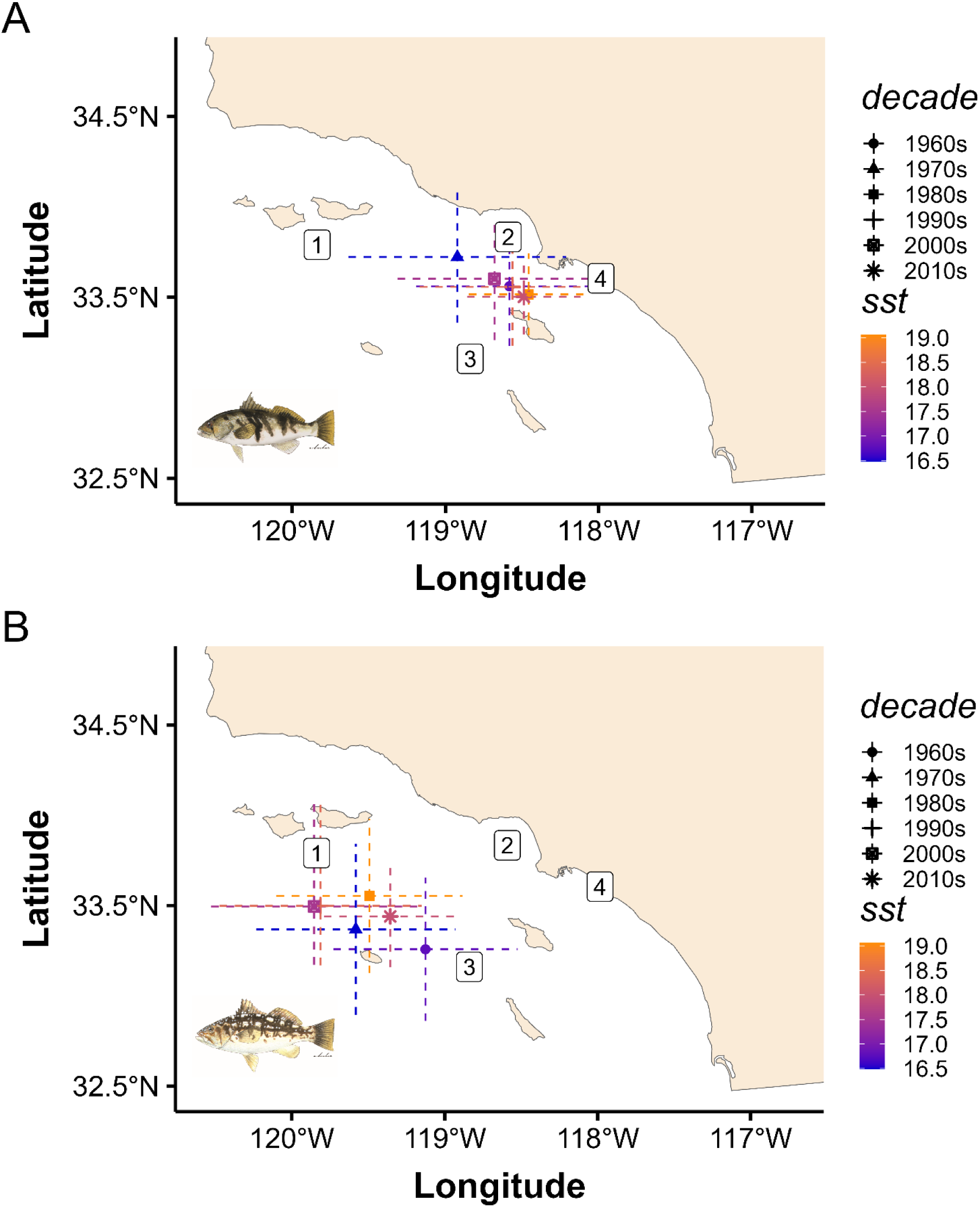
Decadal trends in the mean center of gravity for larval distributions of a) Barred Sand Bass *Paralabrax nebulifer* and b) and Kelp Bass *P. clathratus* in southern California, USA, 1963 to 2016, by decade (symbols) and annual mean sea surface temperature (colors). Lines depict latitudinal and longitudinal 75% confidence intervals. 1 = northern Channel Islands, 2 = Santa Monica Bay, 3 = southern Channel Islands, 4 = Huntington Beach

## Discussion

Our reconstructed larval indices of abundance span over half a century and represent the longest species-specific, fishery-independent time series for the saltwater basses in southern California, USA. Though larval abundance is generally considered to be a suitable proxy for adult spawning stock biomass (Hsieh et al. 2005, 2006), our results indicate southern California bass larvae abundance over the last several decades is a better reflection of future catch in the fishery, as both BSB and KB larval abundance had moderate to strong predictive power in forecasting fish catches. In addition, for both species, this result was consistent across different sources of catch data, i.e., harvest reported in CPFV logbooks, and estimates of total catch (includes harvest and releases across all fishing modes). Moreover, the utility of this predictive power for fishery management is bolstered by the strong relationships we identified between bass larval abundance and environmental variables (e.g., temperature, zooplankton biomass, MLD), and thus, paints a path forward for an ecosystem approach to managing this fishery. Additionally, our analysis revealed species-specific temporal and spatiotemporal trends, as well as differences in the strength of environment-species relationships, suggesting potential differences in their resilience and recovery potential with respect to intrinsic oceanographic variability and anthropogenic climate change impacts.

### Saltwater bass larvae predict future catch

Once considered the “recruitment problem” in fisheries (Haltuch et al. 2019), variability in fishery year-class strength is now understood to be driven by a multitude of factors affecting both pre-and post-recruitment stages, including maternal effects, and is likely to vary by species and population (Houde 2008). Thus, although the challenge of forecasting fishery recruitment remains, research that identifies even modest prediction capabilities is recognized as valuable for informing management. This recognition allows us to more freely accept that prediction may be easier for some species than others, even if we cannot identify all the mechanisms driving those relationships. The added challenge, however, is that predictive ability has been shown to break down for some species due to non-stationary environment-species relationships (White et al. 2019, Litzow et al. 2020), making long-term monitoring and re-evaluation an essential part of these respective research efforts. This is especially critical when environment-species relationships are identified based on shorter time series, as change may include both magnitude and direction. A strength of this study is that our results are based on several multidecadal time series. Thus, at least part of the variability in the relationships we identified between larvae and environmental covariates reflects any inherent long-term interannual and decadal variability influencing the strength of those relationships through time.

In the absence of data on spawning stock biomass, very early life stages like eggs and preflexion larvae, are often a suitable proxy (Hilborn & Walters 1992, Gunderson 1993), such that changes in early larval abundance should reflect changes in female spawner biomass and subsequent recruitment of juveniles. Unfortunately, due to additional, variable mortality after the larval stage, spawning biomass and larval abundance typically show weak relationships with subsequent juvenile and fishery recruitment (Cury et al. 2014, Szuwalski et al. 2015). Our findings with the basses are therefore unusual as we found that young larval abundance was a better predictor of future fish catches than SSB. A strong relationship between spawning stock biomass and larval abundance in the same year relies on 1) appreciable spawning occurring every year, 2) spatiotemporally consistent mortality prior to larvae being surveyed, and 3) that larvae are locally sourced within the region. Thus, in the case of saltwater basses, the breakdown in this relationship may be partly driven by variable larval mortality or variable spawning behavior associated with environmental conditions and harvest impacts. For BSB, there also appears to be a lack of consistent, appreciable, locally sourced larvae from year to year.

We assume the positive relationship between saltwater bass early larval abundance and future fish catches is associated with a survival advantage through adulthood, perhaps imparted by fast initial larval growth and a short preflexion larval stage duration (Fontes et al. 2011, Robert et al. 2023). Though not commonly observed, there are examples when fishery recruitment is set in the first year of life, such that changes in early life stages reflect changes in future fishery year-class strength (Cushing 1990, Stige et al. 2013, White et al. 2019, Schilling et al. 2022). Our findings may also partially be a function of the geographic location of the study. Southern California represents the northern extent of the geographic range for these species, especially for BSB, and populations at their geographic margins tend to experience higher population recruitment variability or recruitment limitation (Myers 1991, Neill et al. 1994, Levin et al. 1997), which is typically reflected in subsequent population size (Armsworth 2002). Schilling et al. (2020), for example, found a positive correlation between predicted Bluefish (Pomatomus saltatrix) larval “settlement” (transformation to the juvenile pelagic phase) and fishery catch-per-unit-effort at the southern end of its distribution.

Regardless of the mechanism, the moderate to strong correlations between bass larval abundance and future catch were observed across multiple catch sets and corresponded to biologically meaningful lags. Both species grow at similar rates, becoming susceptible to hook-and-line fishing gear at two to three years of age and reaching size at fishery recruitment between five and seven years for most of the time series (e.g., in2013, the minimum size limit increased from 12 to 14 inches, which corresponds with fish ages seven to nine years old; Love et al. 1996, Jarvis et al. 2014a, Walker et al. 2020b). Interestingly, when catch including releases was considered, we saw that the species known for a higher catch-and-release rate (KB) showed significant correlations across many more lags (from 3-to 10-y lags) compared to BSB (only 6-to 7-y lags). The highest correlations for both species across both catch data sets corresponded to the age of fishery recruits, while the significant correlations across other lags are likely an artifact of the catch data comprising many cohorts. Thus, restricting the analysis to catch comprising fish at fishery recruitment size or just those fish representing the modal length of the catch may result in even stronger relationships at just one or two lags (Jarvis et al. 2014a, Schilling et al. 2022). We chose not to focus on specific length frequencies because length data were only available for one catch data set, and we wanted to have the ability to test the larva/catch relationship across all available catch data sets.

One potential caveat to using the catch data as a proxy for spawning stock biomass, in the case of BSB, is that hyperstable catches, i.e., stable, high catches, in aggregation-based fisheries can occur during a population decline, resulting in a nonlinear relationship between catch and abundance and an overestimate of abundance at low population sizes (Sadovy de Mitcheson 2016). However, a strong positive linear correlation exists between BSB harvest and fishery-independent adult densities in southern California (Jarvis Mason et al. 2024); thus, there is low probability our results were overly influenced by hyperstability. Moreover, our findings are consistent with other studies showing positive relationships between early life history indices of bass abundance (larvae, young-of-the-year [YOY] juveniles) and future saltwater bass fishery recruitment, despite different collection methods and study time frames (Jarvis Mason et al. 2024, Jarvis et al. 2014a, Miller & Erisman 2014).

### Species-specific differences in bass larval abundance trends

Trends in the standardized indices of larval abundance for both species suggest that the populations of KB and BSB in southern California experienced similar sporadic, strong larval pulses. However, our results also indicate that the KB population has more reliably persisted, being more abundant overall and having less variable and higher larval abundance between strong sporadic recruitment pulses than BSB. This pattern for KB is consistent with locally sourced larval recruitment (Selkoe et al. 2007) and higher densities (Warner 1985). In contrast, BSB larval trends may be more influenced by occasional seeding from Baja California, Mexico (Jarvis Mason et al. 2024), a region we found to have relatively higher BSB larval abundance. Arafeh-Dalmau et al. (2023) demonstrated northward transboundary larval connectivity of up to 500 km between northern Baja California and the SCB during the fall for California Sheephead (*Semicossyphus pulcher*), a species with a slightly longer pelagic larval duration (PLD; 42 days in Cowen 1991) than that of BSB (∼ one month, Findlay & Allen 2002).

The inferred difference in population persistence between the two saltwater basses is also consistent with previous anecdotal reports suggesting BSB has fluctuated in its contribution to the saltwater bass catch, depending on temperature-driven availability (Young 1969, Frey 1971, Feder et al. 1974). Natural log larval abundance trends also identified species-specific differences in decadal-scale availability, with BSB larvae being relatively more abundant during the 1980s and 1990s (a warm period) and KB larvae showing a dramatic, sustained decline during that same period. Nevertheless, larval abundance for both species showed dramatic increases at the end of the time series, which was one of the warmest periods on record (Jacox et al. 2018b). Our results suggest these trends are due to different environmental influences on saltwater bass larvae and different sensitivities to the same environmental influence.

### Temperature, zooplankton, and mixed layer depth predict bass larval abundance

We found strong relationships between saltwater bass larval abundance and temperature, zooplankton biomass, and MLD (here, MLD equals ILD). In general, our model results indicated bass larval abundance was higher when temperatures were warmer, zooplankton biomass was higher, and the MLD was shallower. The relationships between larvae and temperature and zooplankton biomass are not surprising, as peak spawning for KB and BSB occurs during the warm summer months (Erisman & Allen 2005, Jarvis et al. 2014b) following the spring transition in the SCB (McClatchie 2014). Our metric for zooplankton biomass was displacement volume, which can be a crude index of zooplankton community structure as it is highly influenced by pelagic tunicates (e.g., salps) and other large gelatinous organisms (Lavaniegos & Ohman 2007). To better elucidate the potential impact of pelagic invertebrate prey, we explored the relationship between bass larval abundance and biomass of *C. pacificus* from spring cruises (this data was not available for July cruises). We reasoned that increased copepod biomass available to important adult bass prey species (e.g., coastal forage fishes) in the spring might translate into positive maternal conditioning of larvae in summer (Walsh 2023). Inclusion of *C. pacificus*, however, did not improve model performance. Further research is necessary to understand the importance of bass larval prey and maternal effects on bass larval survival.

The MLD had the strongest environmental relationship with bass larval abundance, i.e., shallower MLDs predicted higher bass larval abundance for both species. This relationship might be explained by “seasonal trophic amplification” in nutrient-rich regions like the California Current Ecosystem (CCE; Xue et al. 2022). In these regions, phytoplankton and zooplankton become more concentrated within shallower MLDs, thereby increasing prey encounter rates and, consequently, grazing rates of zooplankton, such that higher feeding efficiency in these regions is driven more by seasonal changes in the depth of the mixed layer than by the amount of food present (Xue et al. 2022). In an analysis of environmental covariates on the biological response of larval fish communities in the CCE, the ILD had the highest ability to predict ecosystem state within the southern California region (Hunsicker et al. 2022). Despite the saltwater basses showing relationships with the same environmental variables, relationships with temperature and zooplankton biomass were relatively stronger for BSB than they were for KB, suggesting BSB is more sensitive to cooler temperatures and declines in zooplankton biomass. Indeed, BSB was historically considered a southern, subtropical/tropical species with an impermanent southern California presence (Young 1963, 1969, Frey 1971, Feder et al. 1974).

If waters off southern California continue to warm, we might expect a shallowing of the MLD due to increased stratification of warm surface waters and thus, higher, less variable, BSB larval recruitment. However, the assumption that climate-driven increased stratification will result in MLD shoaling belies evidence to the contrary (Somavilla et al. 2017), including a recent global analysis that documented a deepening of the MLD over the last five decades at a rate of 5-10 m dec^-1^, despite concomitant increases in stratification (Sallée et al. 2021). Thus, if this trend continues, and southern California saltwater bass abundance is truly a function of the MLD, then a deeper mixed layer might counteract surface warming benefits by decreasing the concentration of prey items. This deepening also has the potential to result in shallow, nearshore depths (“kelp-forest depths”) becoming nutrient-poor (Parnell et al. 2010), which could have negative impacts on KB larval settlement (White and Caselle 2008).

Additional abiotic processes (beyond those examined here) are likely to influence KB larval recruitment dynamics. For example, the timing, direction, and strength of ocean currents impacting larval dispersal and connectivity influences if and where they are transported onshore to settle, particularly in the physically dynamic SCB (Warner 1985). The primary KB larval hotspot was the northern Channel Islands, which is influenced by the “transition zone” in the SCB where many ocean currents converge (Hamilton et al. 2010). White & Caselle (2008) reported that in southern California, the density of Giant Kelp had a positive effect on KB larval settlement, but the relationship was conditioned on larval supply, whereby adult densities were largely a function of whether there was a match between areas of high larval supply and areas of higher kelp stipe density. In that study, the authors explain how eddies and temperature fronts may drive a match-mismatch between KB larval supply and kelp spore supply. This may partially explain why we found that Bight-wide kelp canopy extent was not an important driver of KB larval abundance.

Larval dispersal and retention are likely equally important for BSB. Nearshore flow in the SCB is mostly alongshore with a minor cross-shelf component (Winant & Bratkovich 1981, as cited in Barnett et al. 1984). Whereas we found the predicted geographic distribution of BSB larvae were almost exclusively distributed nearshore along the mainland coast, KB larvae through time showed broad distribution within the SCB with hotspots at the northern Channel Islands, suggesting different processes acting on their distributions (White et al. 2019). Indeed, we observed differences in the two species’ model’s spatiotemporal fields over time that indicate different latent forcing of dynamic biotic and abiotic processes. It is therefore possible that different environmental conditions cue broad scale synchronized spawning events to take advantage of tides and circulation patterns that help retain larvae close to shore (BSB; Barnett et al. 1984) or provide longshore and offshore dispersal to increase the probability of reaching areas with Giant Kelp, including at offshore islands (KB; see asymmetric directionality of KB larval transport in Watson et al. 2010). Differences in spawning aggregation formation and migration may also contribute to spatiotemporal variability in these species’ larval distributions, as BSB form large spawning aggregations at a few predictable locations, while KB spawning aggregations are smaller and more broadly distributed (Erisman & Allen 2006, Jarvis et al. 2010).

All else equal, warmer temperatures for summer spawners seem to be better for higher growth and survival of eggs and larvae (Gadomski & Caddell 1996); however, during the warm regime of the 1980s and 1990s, the SCB shifted to nutrient-poor conditions and experienced a higher frequency of storms, both of which negatively impacted kelp forests in the region (Parnell et al. 2010). Thus, the steady decline in KB larval recruitment during that period may be indirectly associated with decadal-scale declines in Giant Kelp (rather than interannual variability in regional canopy area), despite more optimal temperatures for larval growth and survival (Jarvis et al. 2014a). This could be one driver explaining why our models revealed the influence of temperature on larval KB larval abundance was not as strong as that for BSB.

Although both species showed positive trends with temperature, we did not find evidence of a northward latitudinal shift in the center of larval distribution through time. Although climate-driven phenological and distribution shifts have been well-documented around the world (Pinsky et al. 2020) and historical bass populations extended much farther north than southern California during the region’s tropicalization of the mid-1800s (Hubbs 1948), previous research on larval distribution/phenology shifts in the CCE showed that species responses to climate change are not omnipresent (Hsieh et al. 2009, Asch 2015, Auth et al. 2018, Thompson et al. 2022). In addition, northward larval dispersal of nearshore species beyond the well-known geographic break at Pt. Conception in the SCB, at least during the last sixty years, is thought to be limited more so by hydrographic features than temperature (Warner 1985).

### Implications for fishery recovery and management

Despite the importance of the basses to southern California marine anglers, currently no formal stock assessment exists for either species. Our standardized indices of bass larval abundance offer the potential to develop a species-specific stock assessment model in which the larval abundance index is used as a pre-recruit index or as a means to ground truth an assessment’s predicted recruitment deviations. Even without a stock assessment, our indices offer predictive ability and insight into the potential for each species’ fishery recovery. For example, despite sustained low fishery catches in the early-to-mid-2010s, we found that larval abundance for both species peaked during this time. Given our findings that larval abundance predicts future catches, the increase in larval abundance suggests fishery recovery is imminent. Indeed, fishery-independent indices of adult BSB and KB densities show an upward trend in recent years (CDFW 2021, 2023). Based on larvae/catch relationships reported here, the first peak in recent larval recruitment (2012) should be evident in catches as early as 2020, corresponding to the average fishery recruitment age of eight years. Indeed, recent harvest reports show increases in harvest in 2020 (KB), 2021 (BSB), and 2023 (BSB; PSMFC 2024). In fact, although still relatively low compared to historical harvest, BSB harvest in 2023 was the highest in over a decade, surpassing KB harvest for the first time during the same period (PSMFC 2024).

Collectively, our results pave the way for an ecosystem approach to management of this fishery and bring us closer toward readying the fishery for climate change. Such an approach should consider (among other factors) species-specific differences in larvae-environment relationships and factors influencing differences in temporal and spatiotemporal trends in larval abundance and distributions.

Perhaps one of the most important conclusions of our study is that these two basses have different population dynamics: 1) larval recruitment patterns indicate differences in the persistence of their populations in southern California (e.g., background recruitment levels for KB are higher and less episodic, which is consistent with locally-sourced recruitment (Selkoe et al. 2007) and higher densities (Warner 1985) and has implications regarding their resilience to harvest impacts and climate change), 2) relative to KB, BSB larval recruitment is more closely tied to temperature, which is consistent with historical reports of BSB availability in southern California (Jarvis Mason et al. 2024) and which also has implications for resilience to harvest impacts and climate change, and 3) the offshore islands in the northern SCB are a notable hotspot for KB larval recruitment, which has implications associated with habitat protection (e.g., Marine Protected Areas, Hopf et al. 2022). In contrast, BSB larval recruitment is limited to the mainland coast and given its sporadic nature and more southerly larval distribution in the SCB, may be influenced more so by El Niño-driven sporadic northward advection of larvae from Baja California (Jarvis Mason et al. 2024) as has been suggested for other fishery species in the region (Smith & Moser 1988, Allen & Franklin 1992, Ben-Aderet et al. 2020). This too has implications with respect to harvest impacts and climate change, as future transboundary connectivity between the SCB and Mexico is predicted to be significantly reduced (assuming decreased PLDs and lower kelp persistence), thus highlighting the importance of protecting habitats important for larval connectivity and binational conservation efforts to manage the collective resource (Arafeh-Dalmau et al. 2023).

Environment-species relationships are typically nonstationary and may not only change in magnitude, but direction (White et al. 2019). Here, we had the opportunity to test whether strong environmental relationships we identified using data spanning 54 years (1963-2016) held up using a shorter part of the time series (1984-2016). Although the influence of zooplankton biomass was important in both periods for both species, the influence of temperature on KB larval abundance was weakened/lost in the shorter time series. The shorter time series began just before the well-documented 1988-89 North Pacific climate shift (Hare & Mantua 2000) after which, some species responses to temperature changed (Litzow et al. 2019) and previously established physical (e.g., surface temperature) and ecological relationships with the NPGO (Di Lorenzo et al. 2008) became weaker (Litzow et al. 2020). Thus, we recommend regular monitoring and re-evaluation of the relationships identified here. This recommendation points to the value of maintaining long-term monitoring data streams, like CalCOFI, particularly for other data-limited state-managed and small-scale fisheries. Future research into the role that maternal effects and prey type and condition play in bass larval dynamics will likely provide additional, important ecosystem considerations for managing this culturally and economically important recreational-only fishery.

## Acknowledgements

We thank California Cooperative Fisheries Investigations cruise participants, past and present, N. Bowlin, S. Charter, and M. Human for help with accessing archival specimens and data sheets, L. Bulkeley, K. Farno, L. Martz, R. Quaal, A.C. Salazar Sawkins, and V. Tang for help with sorting samples, and L. Bulkeley for help with identifying the formalin preserved *Paralabrax* spp. larvae to species. The BSB and KB illustrations included in some of the figures are by Amadeo Bachar (Studio ABachar).

