## Supplemental Material for "Environment-driven trends in fish larval abundance predict fishery recruitment in two temperate reef congeners: Mechanisms and implications for fishery recovery under a changing ocean"

***S1**. Specification of geostatistical mesh*

This type of spatial model uses a predictive-process stochastic partial differential equation (SPDE) triangulated mesh to approximate the spatial component of the Gaussian random fields via interpolation between the mesh vertices (knots, Anderson et al. 2022). For the standardized index of abundance models, we specified a minimum cutoff distance of eight km and incorporated a physical barrier mesh to account for the California coast and islands, in which we specified a spatial range of 0.1, i.e., fractional distance of spatial independence, which corresponds to a spatial correlation decay rate that is ten times faster over land than over water (Fig. S1, Anderson et al. 2022). The specified spatial resolution of the knots was fine enough to prevent overfitting of the model, given the number of data points. For the environmental models, we specified a slightly larger minimum cutoff distance of ten km.

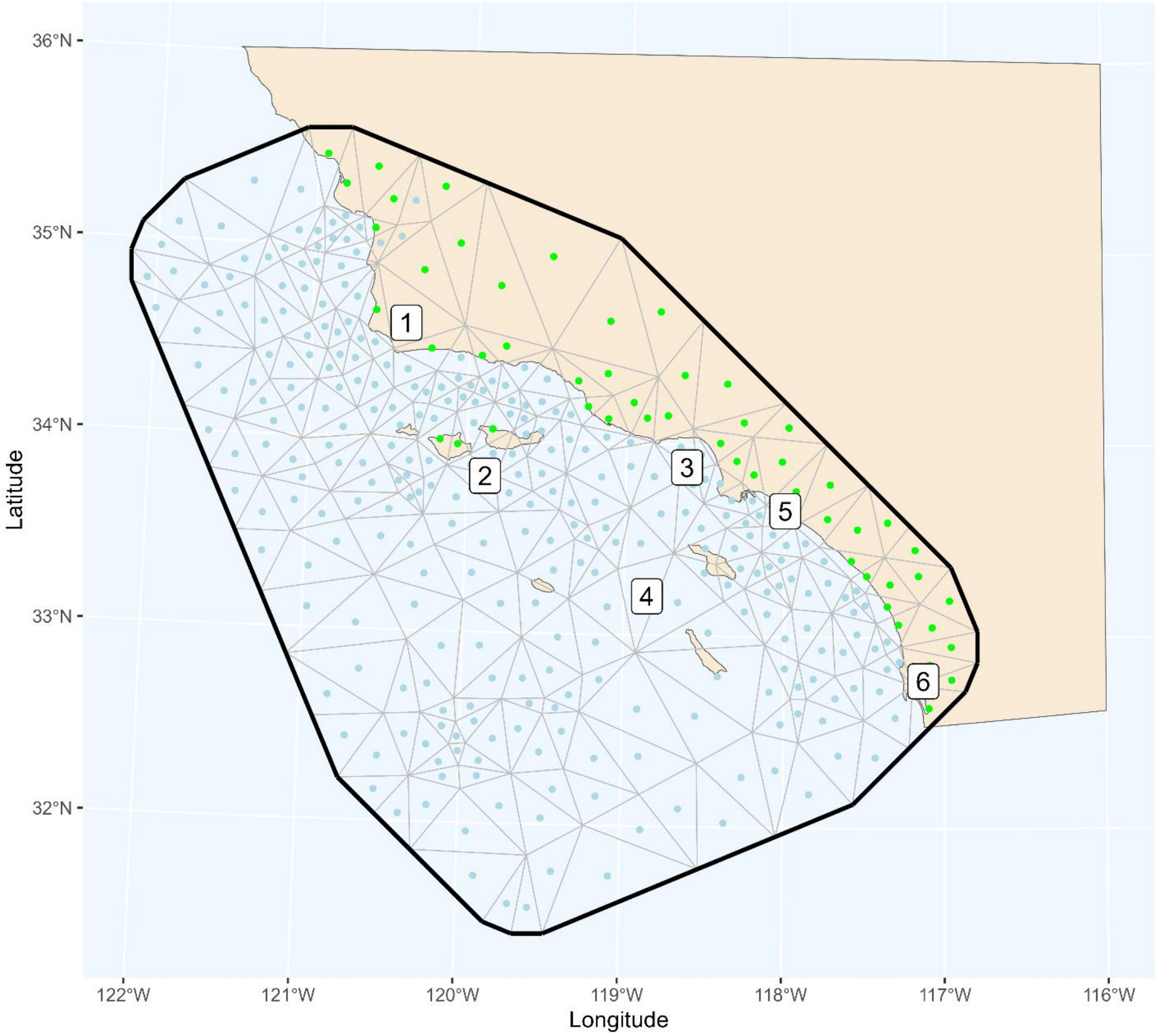


**Figure S1.** Geostatistical mesh (triangles with blue dots) and barrier mesh (triangles with green dots) generated with the R package `sdmTMB` and based on *Paralabrax* spp. larvae data collected in July CalCOFI cruises off southern California, USA, 1963-2016. CalCOFI = California Cooperative Fisheries Research. 1= Pt. Conception, 2 = northern Channel Islands, 3 = Santa Monica Bay, 4 = southern Channel Islands, 5 = Huntington Beach, 6 = San Diego

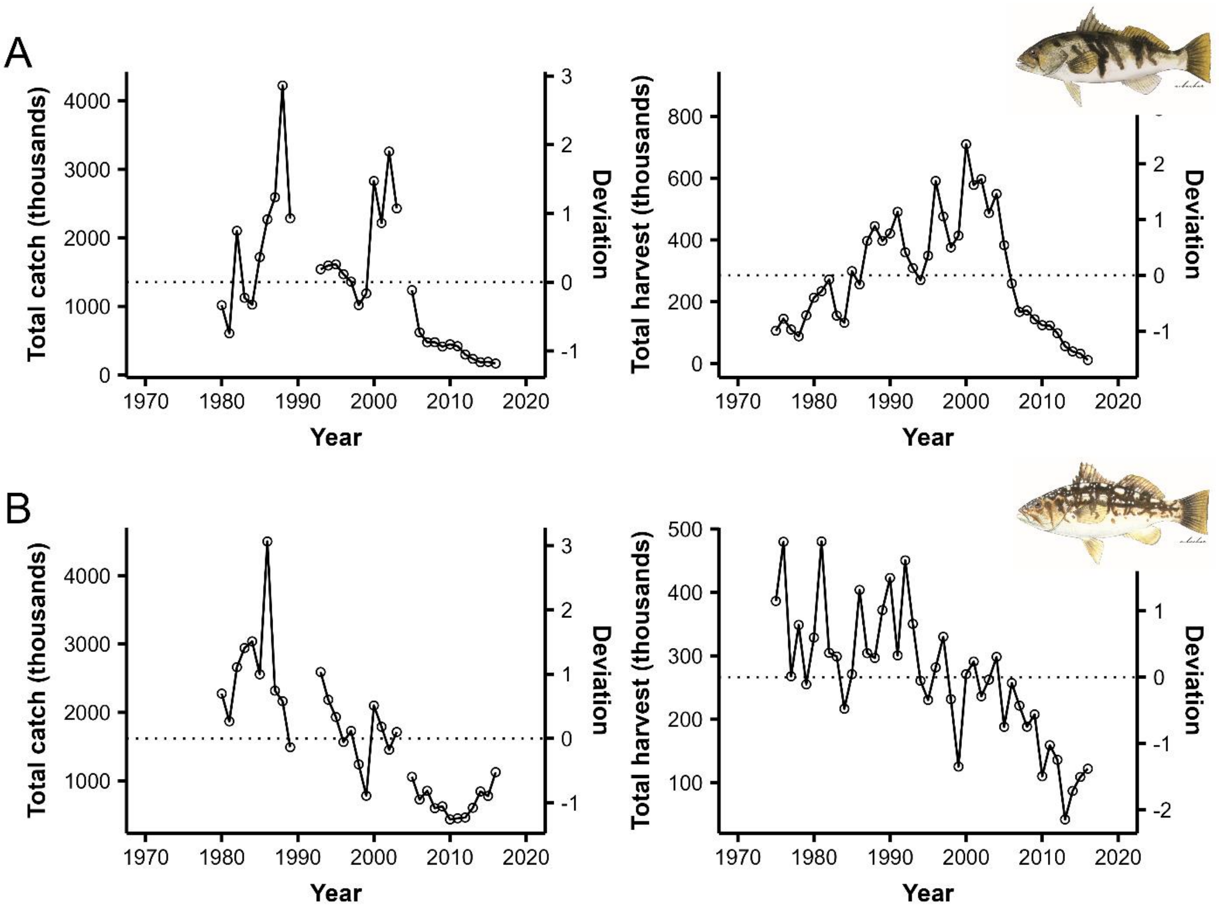


**Fig. S2.** Total estimates of catch (harvest, released alive/dead) across all fishing modes (left) and total reported harvest by Commercial Passenger Fishing Vessels (CPFVs, right) in southern California, USA for a) Barred Sand Bass *Paralabrax nebulifer* and b) and Kelp Bass *P. clathratus*. Total estimates represent data from the Marine Fisheries Statistical Survey (1980-2003) and the California Recreational Fisheries Survey (2005-2016; PSMFC 2024) and CPFV harvest data represent California Department of Fish and Wildlife CPFV logbook data (1975-2016).

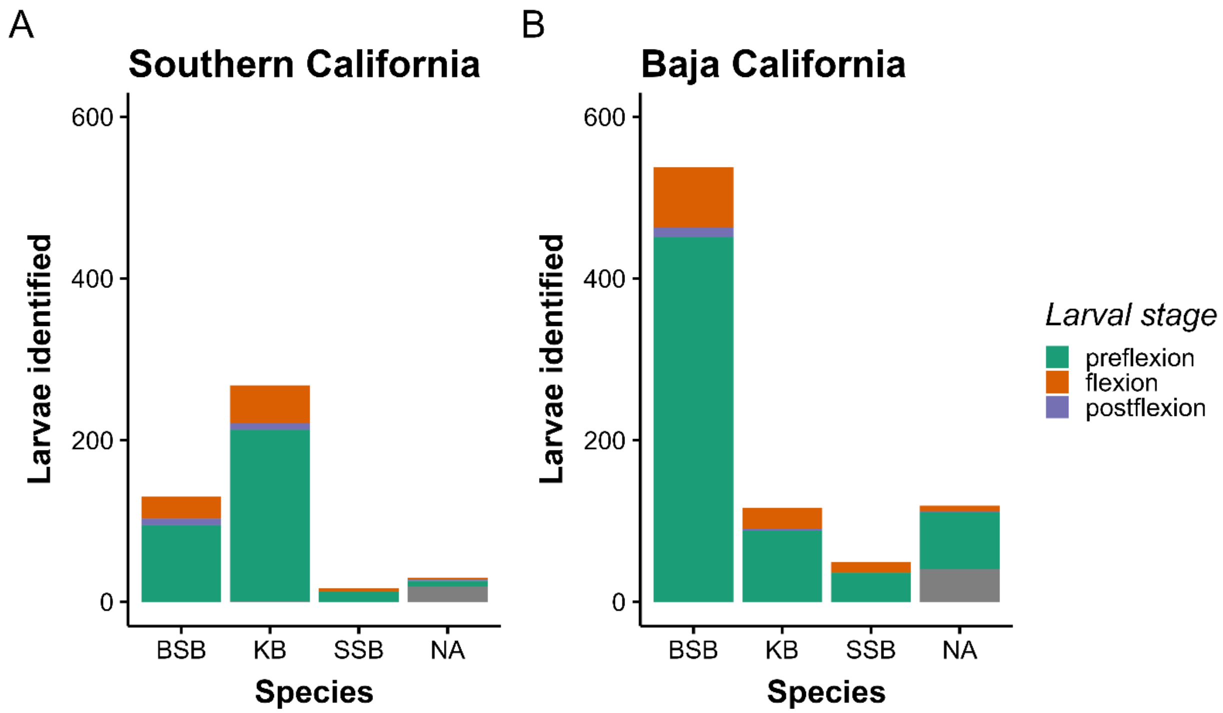


**Figure S3.** Numbers of identified formalin preserved *Paralabrax* spp. larvae, by species and larval stage, originally collected in July CalCOFI cruises off a) southern California, USA, and b) Baja California, Mexico, from 1963 to 2016. CalCOFI = California Cooperative Fisheries Investigations.

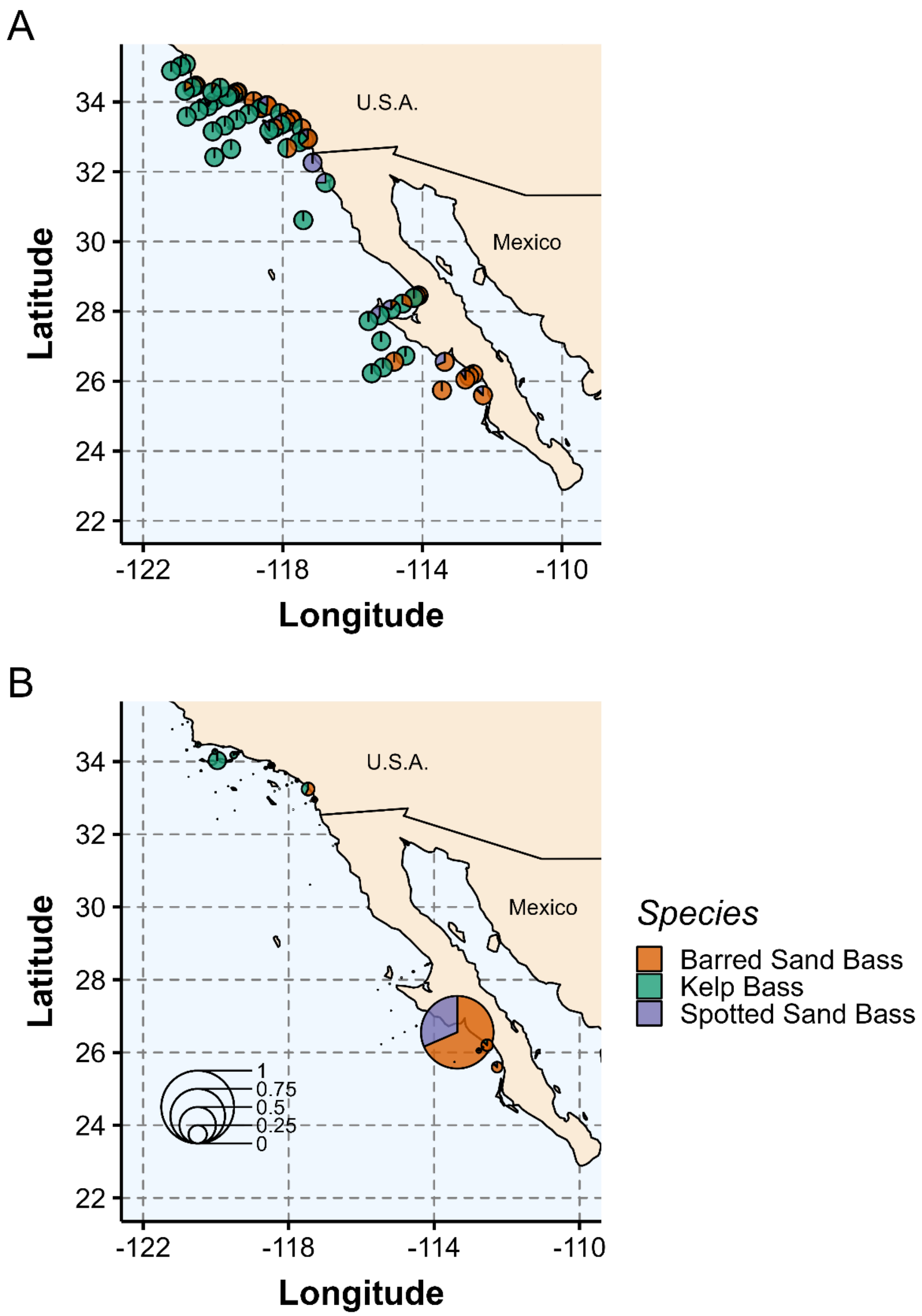


**Figure S4.** Maps of a) the relative proportion of formalin preserved *Paralabrax* spp. larvae identified by species and station from July CalCOFI cruises, and b) the same as (a) above, standardized by the maximum mean number of larvae per species per station (= 12.4 larvae) over the entire survey period from 1963 to 2016. CalCOFI = California Cooperative Fisheries Investigations.

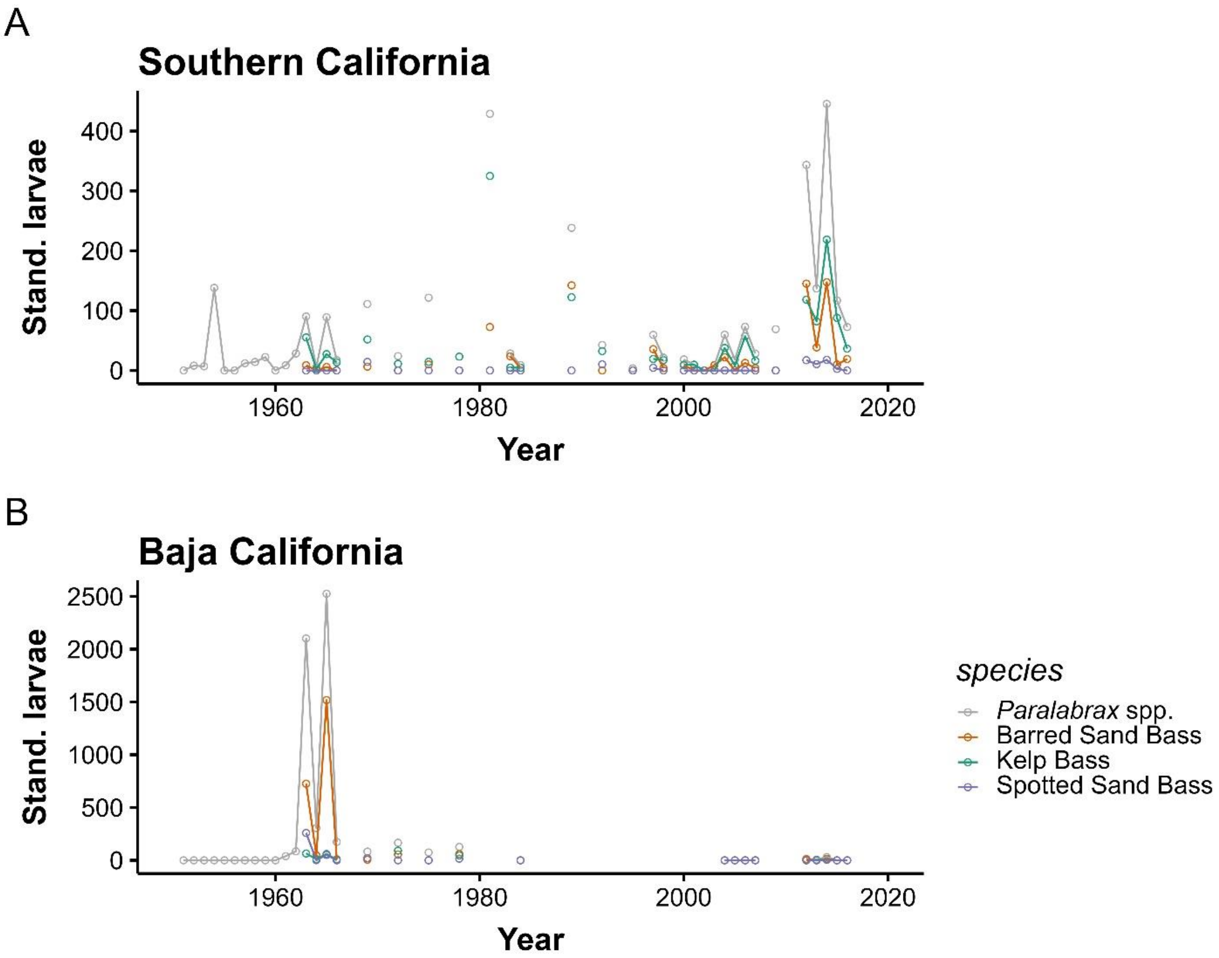


**Figure S5.** Temporal trends in the standardized raw larval counts (larvae under ten $m^2$ of sea surface) for *Paralabrax* spp. larvae combined and by species (Barred Sand Bass *P. nebulifer*, Kelp Bass *P. clathratus,* Spotted Sand Bass *P. maculatofasciatus*) collected in July CalCOFI cruises in a) southern California, USA, and b) Baja California, Mexico, 1950-2016. Note that *Paralabrax* spp. larvae counts in some years may include *P. auroguttatus* (Goldspotted Sand Bass), especially off Baja California. CalCOFI = California Cooperative Fisheries Investigations.

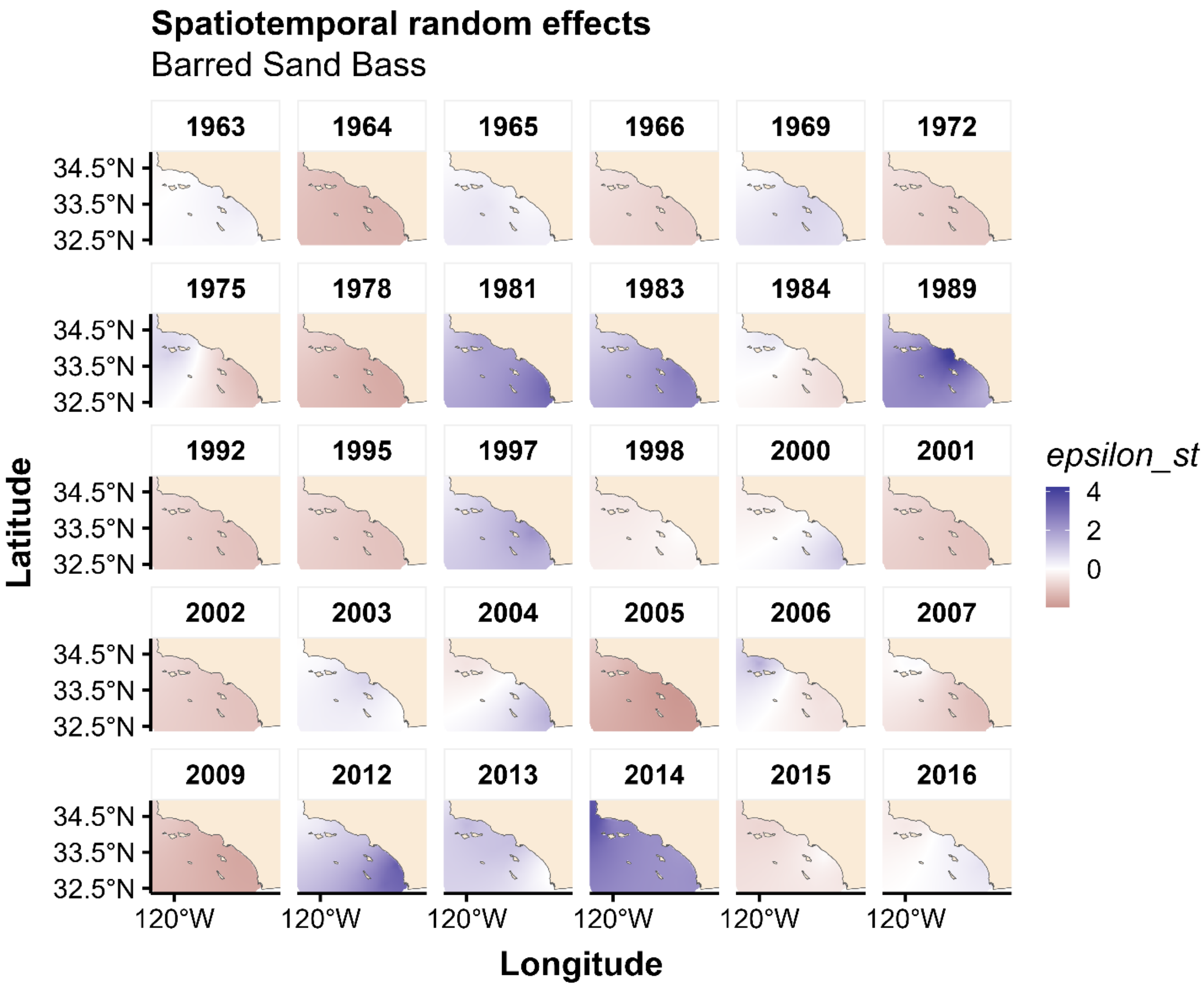


**Figure S6.** Deviations from the fixed effect predictions and spatial random effect deviations derived from the standardized index of abundance model for Barred Sand Bass *Paralabrax nebulifer* in southern California, USA, 1963-2016. Deviations represent the spatiotemporal influence of latent variables on Barred Sand Bass larval abundance, i.e., biotic and abiotic factors that are changing through time and that are not accounted for in the model.

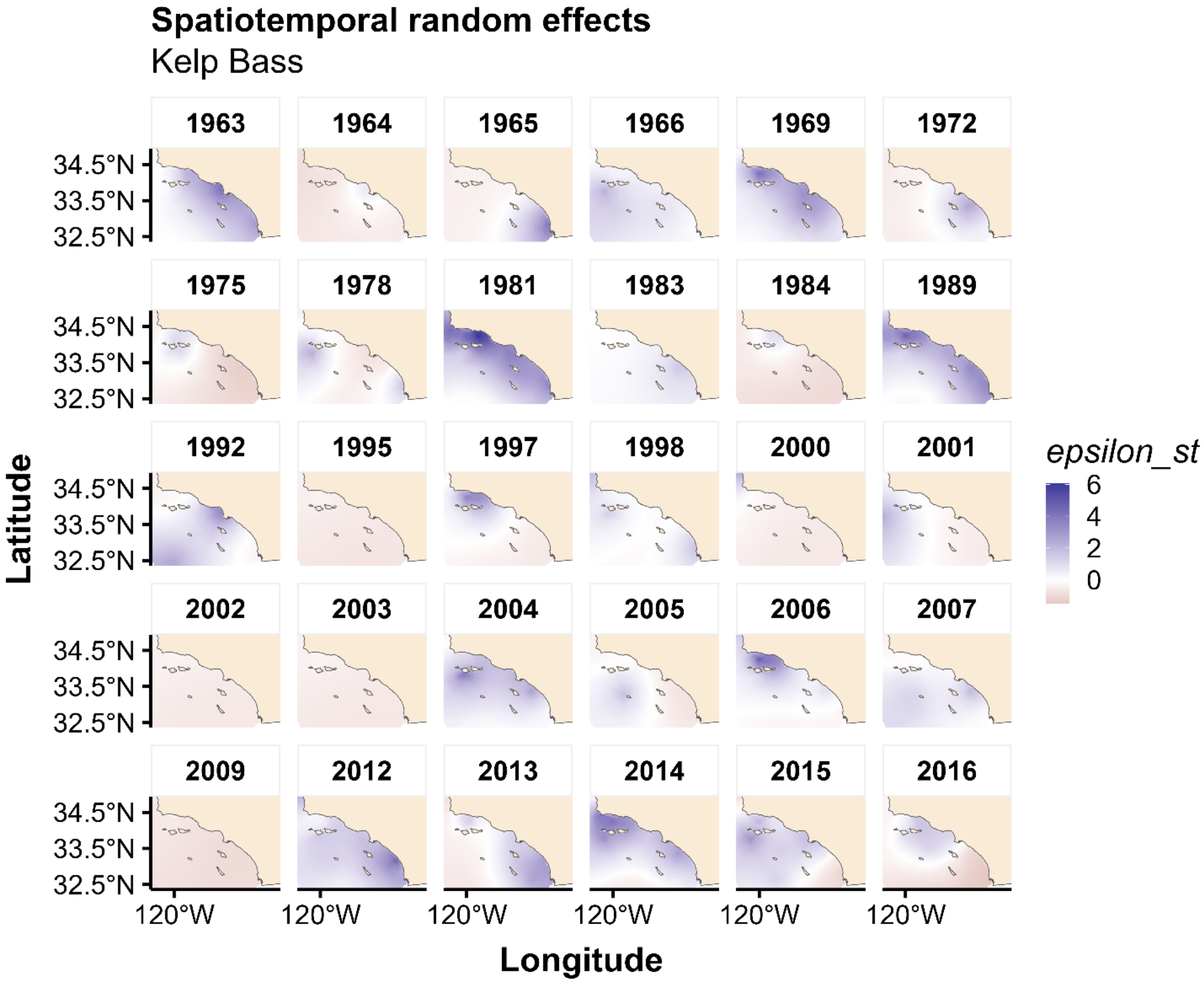


**Figure S7.** Deviations from the fixed effect predictions derived from the standardized index of abundance model for Kelp Bass *Paralabrax clathratus* in southern California, USA, 1963-2016. The model did not contain a spatial random effect. Deviations represent the spatiotemporal influence of latent variables on Kelp Bass larval abundance, i.e., biotic and abiotic factors that are changing through time and that are not accounted for in the model.

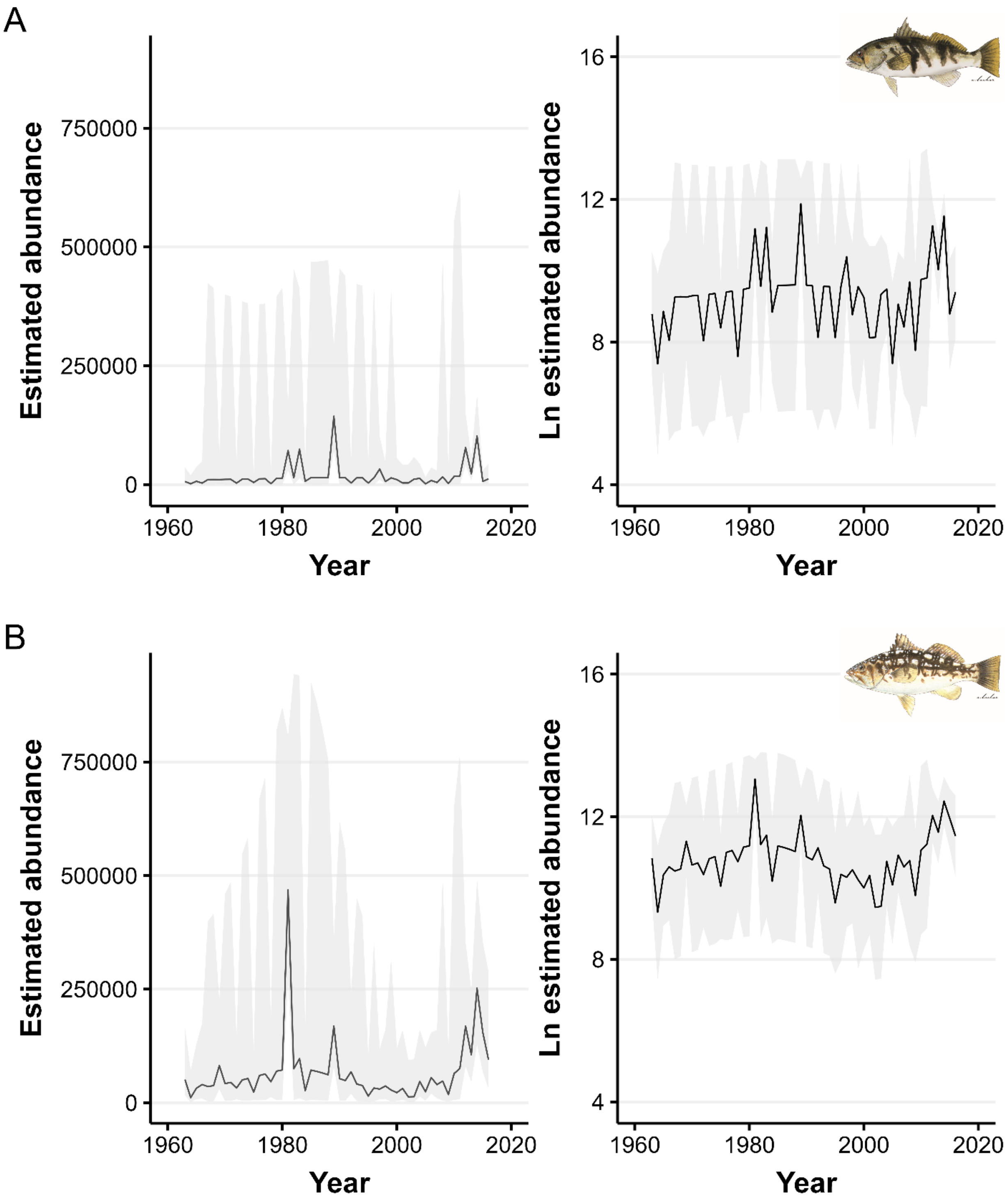


**Figure S8.** Standardized index of larval abundance (link scale, left; natural log scale, right) based on sdmTMB model predictions for all years from 1963 to 2017 (no missing years) for a) Barred Sand Bass *Paralabrax nebulifer* and b) and Kelp Bass *P. clathratus* in southern California, USA. These indices were used for cross correlation analysis with southern California catch data sets.

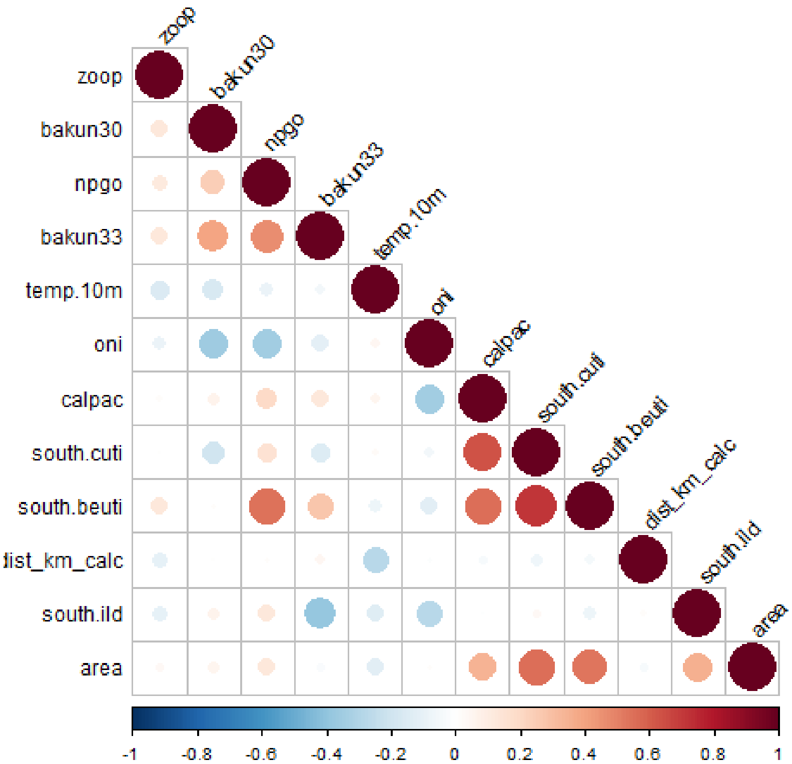


**Figure S9**. Correlation matrix of environmental variables selected for modeling species-specific *Paralabrax* spp. larval abundance in southern California, USA. zoop = CalCOFI zooplankton biomass, bakun30 = Bakun upwelling index at 30°N, npgo = North Pacific Gyre Oscillation index for July, bakun33 = Bakun upwelling index at 33°N, temp.10m = CalCOFI temperature averaged over the upper 10 m, oni = Ocean Niño Index for June/July, calpac = biomass of the calanoid copepod *Calanus pacificus*, south.cuti = Coastal Upwelling Transport Index. south.beuti = Biologically Enhanced Upwelling Index, dist_km_calc = distance to mainland coast, south.ild = isothermal layer depth, area = areal canopy extent of Giant Kelp *Macrocystis pyrifera.* CalCOFI = California Cooperative Fisheries Investigations.

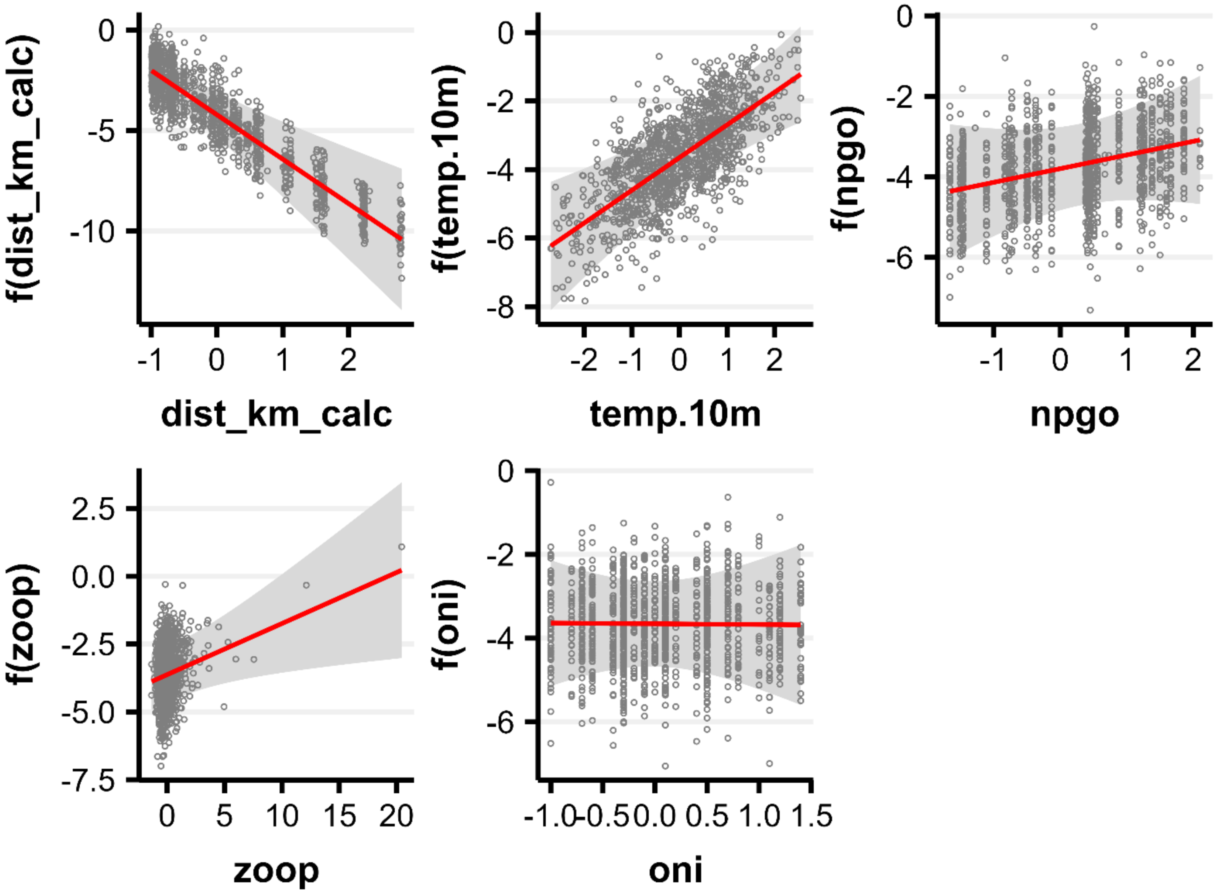


**Figure S10.** Model 1 conditional effects plots representing the predicted effects of select environmental covariates on Barred Sand Bass *Paralabrax nebulifer* larval abundance in southern California, USA, 1963-2016. Shaded ribbon depicts the 95% confidence band, and the points depict partial residuals. Individual covariate effects are conditioned on the other covariates being fixed at their median values. dist_km = distance to mainland coast, zoop = CalCOFI zooplankton biomass, temp = CalCOFI temperature averaged over the upper 10 m, oni = Ocean Niño Index for June/July, npgo = North Pacific Gyre Oscillation index for July. CalCOFI = California Cooperative Fisheries Investigations.

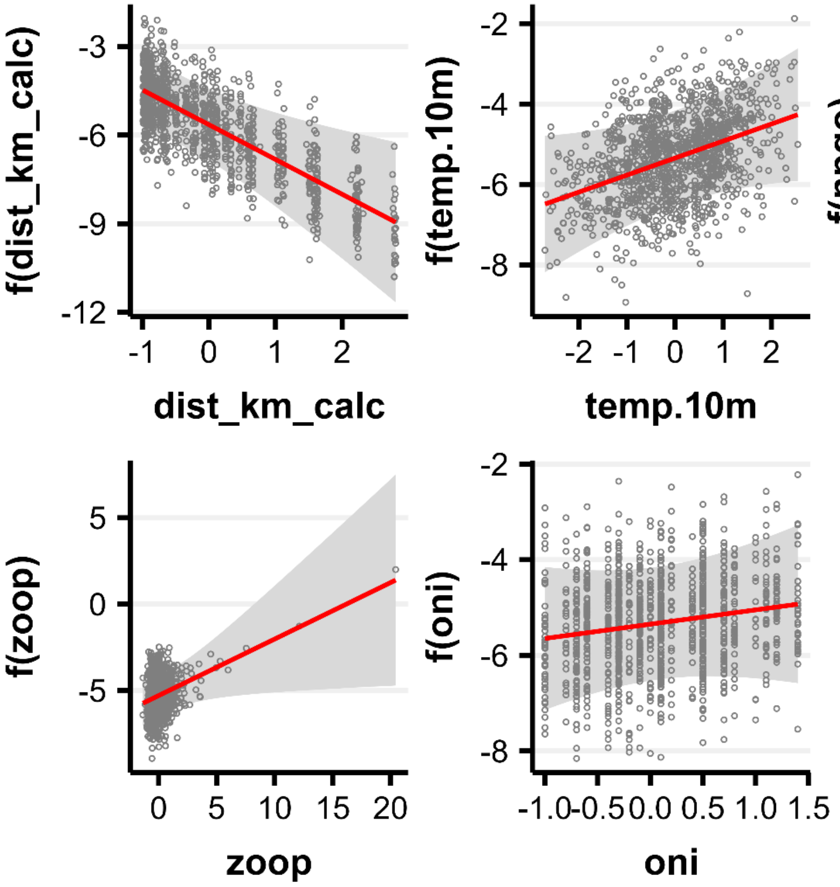


**Figure S11.** Model 1 conditional effects plots representing the predicted effects of environmental covariates on Kelp Bass *Paralabrax clathratus* larval abundance in southern California, USA, 1963-2016. Shaded ribbon depicts the 95% confidence band, and the points depict partial residuals. Individual covariate effects are conditioned on the other covariates being fixed at their median values. dist_km = distance to mainland coast, zoop = CalCOFI zooplankton biomass, temp = CalCOFI temperature averaged over the upper 10 m, oni = Ocean Niño Index for June/July, npgo = North Pacific Gyre Oscillation index for July. CalCOFI = California Cooperative Fisheries Investigations.

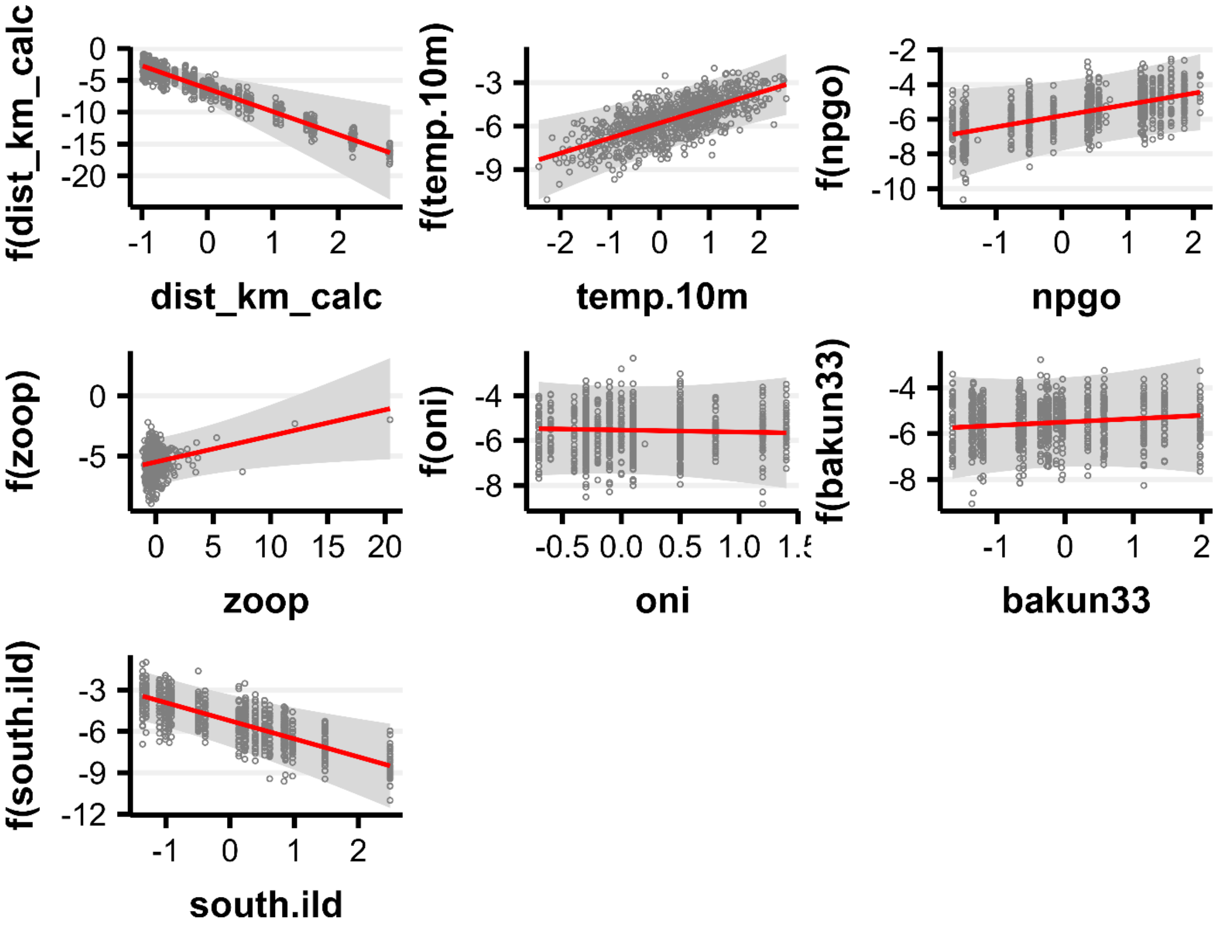


**Figure S12.** Model 2 conditional effects plots representing the predicted effects of environmental covariates on Barred Sand Bass *Paralabrax nebulifer* larval abundance in southern California, USA, 1984-2016. Shaded ribbon depicts the 95% confidence band, and the points depict partial residuals. Individual covariate effects are conditioned on the other covariates being fixed at their median values. dist_km = distance to mainland coast, zoop = CalCOFI zooplankton biomass, south.ild = isothermal layer depth, temp.10m = CalCOFI temperature averaged over the upper 10 m, oni = Ocean Niño Index for June/July, npgo = North Pacific Gyre Oscillation index for July, and bakun33 = Bakun upwelling index at 33°N. CalCOFI = California Cooperative Fisheries Investigations.

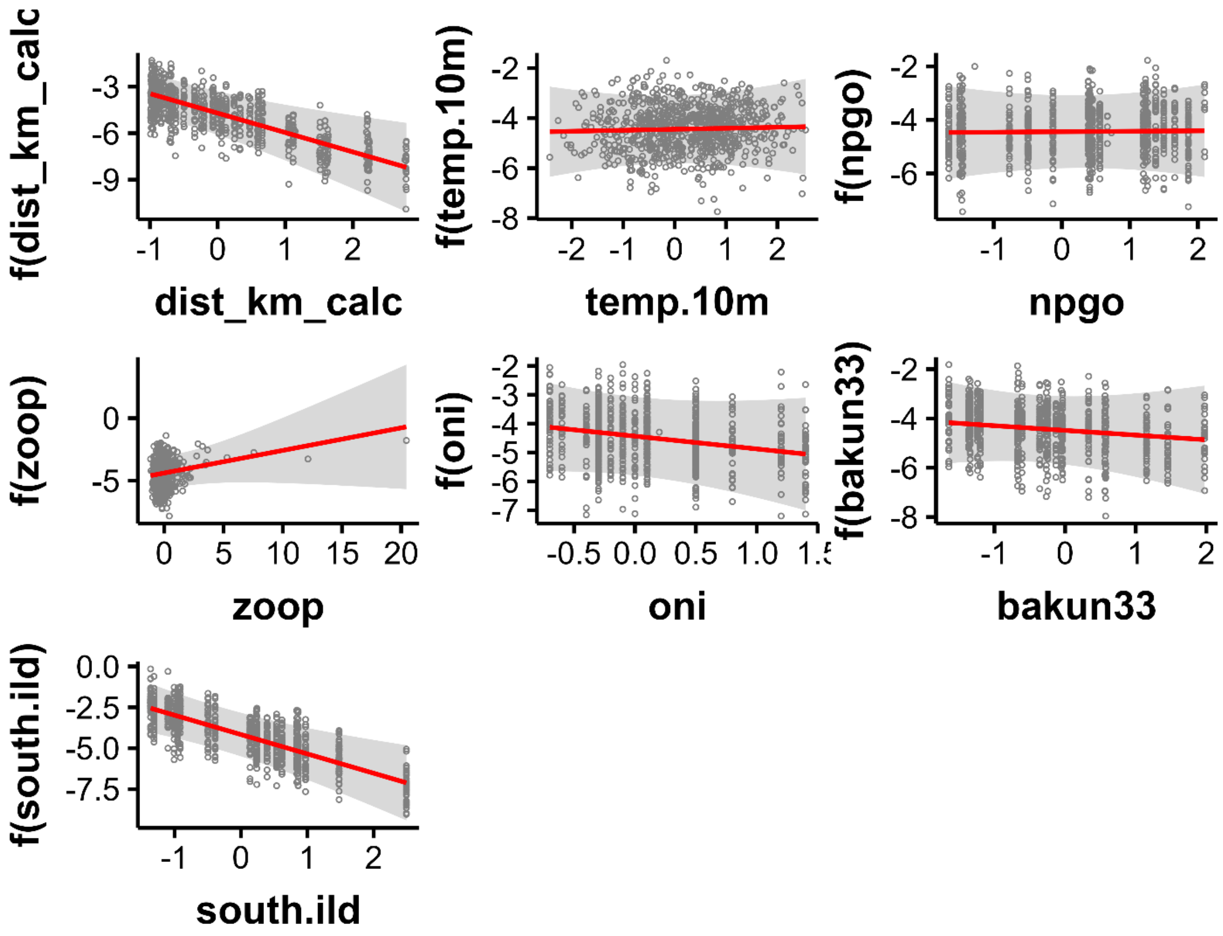


**Figure S13**. Model 2 conditional effects plots representing the predicted effects of environmental covariates on Kelp Bass *Paralabrax clathratus* larval abundance in southern California, USA, 1984-2016. Shaded ribbon depicts the 95% confidence band, and the points depict partial residuals. Individual covariate effects are conditioned on the other covariates being fixed at their median values. dist_km = distance to mainland coast, zoop = CalCOFI zooplankton biomass, south.ild = isothermal layer depth, temp.10m = CalCOFI temperature averaged over the upper 10 m, oni = Ocean Niño Index for June/July, npgo = North Pacific Gyre Oscillation index for July, and bakun33 = Bakun upwelling index at 33°N. CalCOFI = California Cooperative Fisheries Investigations.
